# Actin-based deformations of the nucleus control multiciliated ependymal cell differentiation

**DOI:** 10.1101/2023.10.13.562225

**Authors:** Marianne Basso, Alexia Mahuzier, Syed Kaabir Ali, Anaïs Marty, Marion Faucourt, Ana-Maria Lennon-Duménil, Ayush Srivastava, Michella Khoury Damaa, Alexia Bankole, Alice Meunier, Ayako Yamada, Julie Plastino, Nathalie Spassky, Nathalie Delgehyr

## Abstract

Ependymal cells (ECs) are multi-ciliated cells in the brain that contribute to cerebrospinal fluid flow. ECs are specified at embryonic stages but differentiate later in development. Differentiation depends on genes such as GEMC1 and MCIDAS with E2F4/5, but also on cell cycle related factors. In the mouse brain, we observe that nuclear deformation accompanies EC differentiation. Tampering with these deformations either by decreasing F-actin levels or by severing the link between the nucleus and the actin cytoskeleton blocks differentiation. Conversely, increasing F-actin by knocking out the Arp2/3 complex inhibitor Arpin or artificially deforming the nucleus activates differentiation. These data are consistent with actin polymerization triggering nuclear deformation and jump-starting the signaling that produces ECs. A player in this process is the retinoblastoma1 (RB1) protein whose phosphorylation prompts MCIDAS activation. Overall, this study reveals an important role for actin-based mechanical inputs to the nucleus as controlling factors in cell differentiation.

## INTRODUCTION

Multiciliated Ependymal cells (ECs) are located along brain ventricles. They form a protective barrier to the brain parenchyma and contribute to cerebrospinal fluid flow through the beating of their motile cilia^1^. Impairment of this flow correlates with age-related dementia^2^, schizophrenia^3^ and hydrocephalus^4^, diseases whose onset is not always understood.

In mammalian brain, lineage tracing experiments show that ECs are sister cells of the adult neural stem cell^5,6^, derived from embryonic progenitors^7^, but acquiring their own identity at later stages. Differentiation of multiciliated cells depends on Notch inhibition that is required for the activation of the master regulators GEMC1 followed by MCIDAS and other effectors^8^. Both GEMC1 and MCIDAS interact with E2F4 or 5 and DP1 to promote transcriptional control of multiciliated cell differentiation^9^. Cell cycle related factors, such as Cdk1 and 2, are also involved in the differentiation of these post-mitotic cells^10,11^. The cues that trigger the activation of the different factors required for differentiation are not clear.

Neural stem cell progenitors are sensitive to forces during their proliferative and differentiating stages in neurons or oligodendrocytes^12^, and it is known in stem cells in general that forces can be directly transmitted to the nucleus through the cytoskeleton^13^. Key players in this transmission are the LINC complexes that span the nuclear envelope and link the nucleoskeleton to the cytoskeleton^14^. In these complexes, the inner nuclear membrane proteins, the SUN family, contact the lamina (nucleoskeleton) and interact with a specific domain (KASH domain) of the outer nuclear membrane proteins (the NESPRIN family encoded by the Syne genes). NESPRINS interact with either microtubules, intermediate filaments or actin. Actin has been shown to contribute to nuclear morphology and positioning in different systems (e.g. migrating fibroblasts^15^, oocytes^16^). The LINC complexes and the mechanical forces transmitted by those complexes to the nucleus are involved in many aspects of cellular functions. For example forces participate in the orientation and shape of the nucleus^17,18^, the positioning of the centrosome^19^, DNA damage and repair^20-22^, changes in gene expression^23-26^ and the opening of nuclear pores or channels^27,28^ with consequences for cell cycle^29-33^ or fate choice^34-37^.

In this context, we sought to uncover the cues that drive EC differentiation. We observed a connection between actin accumulation, nuclear deformation and differentiation. Other results imply that RB1 is involved in transducing nuclear deformations to activate EC differentiation.

## RESULTS

### Nuclei deform during EC differentiation

To visualize nuclear shape during EC differentiation, we stained lateral ventricles of mice expressing Centrin2-GFP^38^ during the process of differentiation (Post-natal day 5, P5) with Lamin-B1 antibodies. We observed a large diversity of shapes and localizations of nuclei (Figure 1A). The Centrin2-GFP allowed us to determine the stage of EC differentiation. Cells at the surface of the ventricle with two Centrin dots are progenitors. Then a cloud or ring-like structures of Centrin appear near the two Centrin dots, indicating the formation of pro-centrioles around deuterosomes (differentiating cells). Finally, several dots (on average 50^39^) of Centrin are released from the deuterosomes and migrate toward the plasma membrane (multi basal body cells) where they dock to form the basal bodies at the base of motile cilia^40^. In the tissue, nuclei are quite close to each other, sometimes in depth; it was thus challenging to associate each nucleus with the corresponding Centrin staining. To solve this problem, we electroporated Centrin2-GFP expressing brains at the onset of differentiation (P1) with membrane-cherry (mb-cherry). This reporter stained the membranes of the electroporated cells allowing the identification of which nucleus goes with which apical surface, where centrioles are located, to assess differentiation stages. On electroporated lateral walls stained for Lamin-B1, we noted that during the process of differentiation (P5) the nuclei increase in volume (1.7x increase on average between progenitor and multi-basal body stages, Figure 1A, B), change shape (becoming 2x flatter at the poles, Figure 1A, C) and migrate toward the plasma membrane (moving around 2 μm up on average, Figure 1A, D). Overall nuclei were drastically changing during the process of ependymal differentiation.

**Figure 1:**
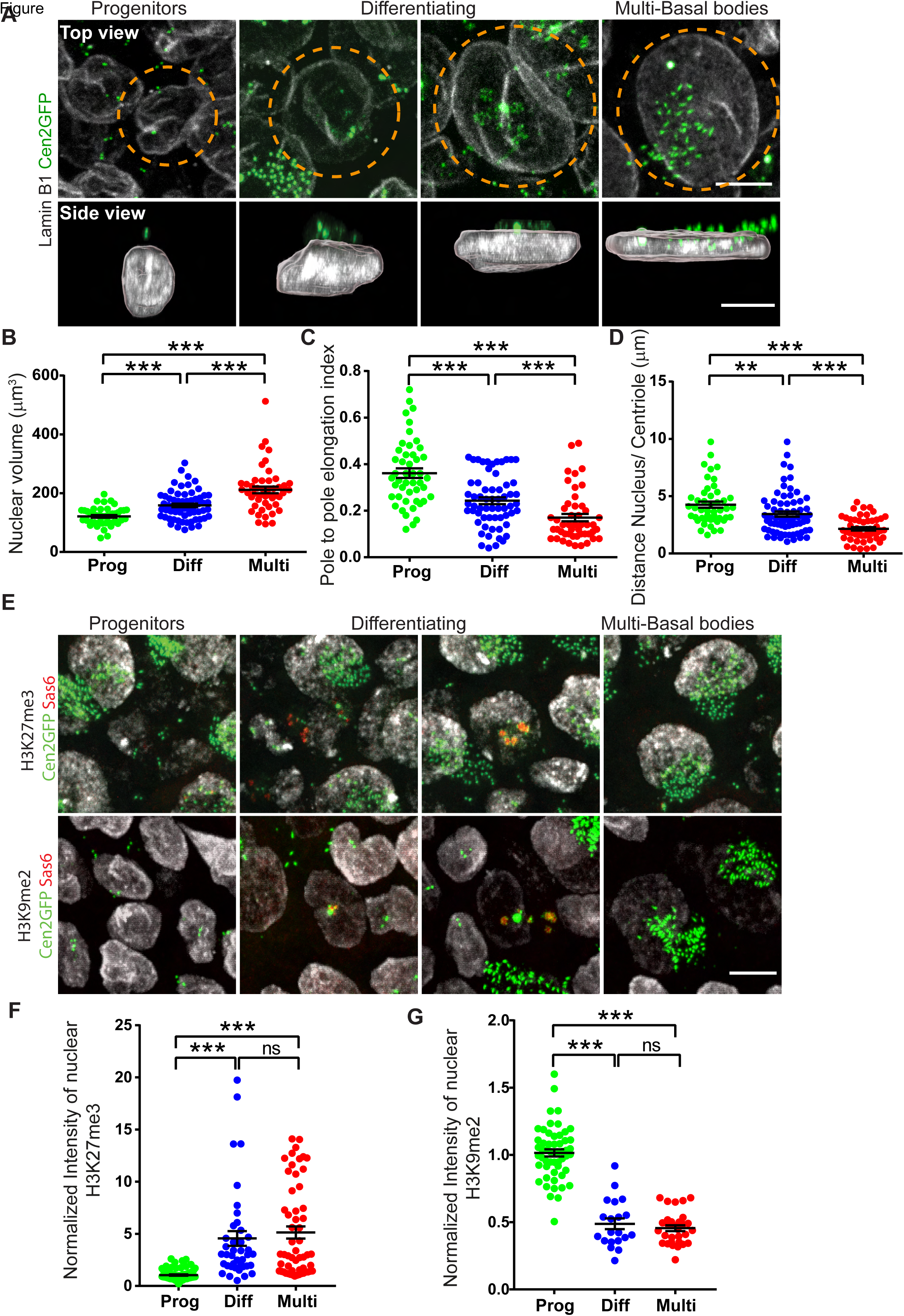
Nuclei deform during ependymal differentiation. All staining were performed on whole mount lateral wall during ependymal differentiation (post-natal day 5, P5). The different stages of differentiation are recognized according to centriolar markers (Centrin=centrioles without **(A)** or with **(E)** Sas6= pro-centrioles): 2 dots of Centrin and no Sas6= Progenitors, 2 dots + clouds or rings of Centrin and dots of Sas6= differentiating progenitors, several dots (more than 5) and no more Sas6 = multi-basal bodies. **A)** Staining of Lamin-B1 (nuclear lamina, Gray) and Centrin2-GFP (Cen2GFP, centrioles, Green) during ependymal cell differentiation (postnatal day 5, P5) on whole mount lateral wall of Centrin2-GFP mice. Top view is a maximum projection of z-stacks. Side view is an Imaris 3D projection of the above cell (orange dashed circle) with segmented nuclei. **(B, D)** Quantification of the nuclear shape during the different steps of ependymal differentiation at P5 on Imaris segmented nuclei of cells transfected at P1 with membrane-cherry (mb-cherry) to be able to associate the apical surface, where the centrioles are located, and the nucleus. **B)** Volume **C)** Pole to pole elongation index (Prolate Ellipticity) **D)** Distance between the center of the nucleus and the highest centriole. Error bars represent the sem of n=54 progenitors (Prog), 66 differentiating (Diff) and 54 multi-basal bodies (Multi) cells from nine mice; p-values were determined by one-way ANOVA followed by Dunn’s multiple comparison test; *** P≤0.001; ** P≤0.01. **E)** Staining of H3K27me3 (Gray, top panel) or H3K9me2 (Gray, bottom panel), Centrin2-GFP (Cen2GFP, centrioles, Green) and Sas6 (pro-centriole, Red) during ependymal differentiation (P5). **(F, G)** Quantification of the fluorescence intensity of H3K27me3 (**F**) or H3K9me2 (**G**) staining in cells expressing mb-cherry during the different steps of ependymal differentiation at P5, normalized to the mean of intensity in surrounding untransfected progenitor cells. Error bars represent the sem of n=76/ 54 progenitors, 40/ 20 differentiating and 56/ 30 multi-basal bodies cells from 3 mice for H3K27me3 and H3K9me2 respectively; p-values were determined by one-way ANOVA followed by Dunn’s multiple comparison test; *** P≤0.001; ns P>0.05. Scale bar = 5 μm.

In the literature, there are reports that mechanical forces applied to the nucleus drive chromatin re-organization and cell fate acquisition ^41,42^. We thus wondered whether the observed changes in nuclear shape and localization were accompanied by a change in chromatin organization. To assess this, we stained Centrin2-GFP mouse lateral wall (electroporated at differentiation onset (P1) with mb-cherry) in the process of ependymal differentiation (P5) with Sas6 (a pro-centriole marker present during the differentiating stage only) and the heterochromatin markers H3K27me3 or H3K9me2. Similarly to the response observed in cells submitted to an artificial mechanical stress^43^, we observed that H3K27me3 increased while H3K9me2 decreased with nuclear re-shaping and differentiation (Figure 1E-G). Thus, nuclei of differentiating cells are actively deforming and re-arranging their chromatin.

### Actin, but not microtubules, are involved in nuclear deformation and EC differentiation

Nuclear deformation and localization in many systems depend on the cell cytoskeleton^44,45^. F-actin and microtubule staining on lateral wall during EC differentiation showed that both networks undergo massive rearrangement, accumulating near the centrioles during the first steps of differentiation (Figure S1A-D). To explore the contribution of both F-actin and microtubules to nuclear shape change, we treated explants of lateral walls containing cells in the process of differentiation (P5) with drugs leading to the depolymerization of microtubules (nocodazole) or actin (cytochalasin D) for 20 hours. We then fixed and stained the explants for Lamin-B1. Cytochalasin-D perturbed the morphology of the nuclei of differentiated cells while nocodazole had no obvious effect (Figure S1E). In keeping with this, interfering with the microtubule molecular motor dynein had no effect on either nuclear deformation (Figure S2A-C) or EC differentiation (Figure S2D-G). The slight effect observed in nuclear position in relation to the centrioles at the progenitor stage might be a consequence of impairment of the interkinetic movement observed in the few remaining dividing progenitor cells at P5 (most of the division happening during embryonic development^7,46^).

Actin cables are enriched at the apical surface of differentiating cells with long cables reaching the nuclei, suggesting that the actin network might cap the nucleus or pull on it. Interestingly, inhibiting myosin activity using blebbistatin treatment for 20 hours on lateral ventricle explants bearing differentiating ECs also affected the deformation of multiciliated cell nuclei (Figure S1E). A host of different actin binding proteins are known to contribute to the actin network in differentiated multiciliated cells^47^. To assess whether F-actin accumulation during differentiation depends on the Arp2/3 complex (branched F-actin) or on Formins (linear F-actin), we treated explants of lateral walls bearing differentiating ECs (P5) with inhibitors of the Arp2/3 complex (CK666) or of Formins (SMIFH2) for 20 hours and then fixed and stained them with phalloidin (Figure S1F, G). Only the treatment with CK666 decreased the accumulation of F-actin in differentiating cells, suggesting a contribution of the Arp2/3 complex to F-actin formation in ECs.

To perturb Arp2/3 complex activity *in vivo*, we electroporated at differentiation onset (P1) an shRNA against *Arp3* for which the sequence was validated^48^. ARP3 is a subunit of the Arp2/3 complex. We performed phalloidin staining at P5 on electroporated lateral walls and showed that sh*Arp3* decreased F-actin accumulation in differentiating cells compared to control and relative to untransfected surrounding progenitor cells (Figure S3A, B). Lamin-B1 staining in cells depleted for *Arp3* revealed that the nuclei do not flatten as much as in control electroporated cells, and migrate less toward the apical side (Figure 2A-C), while nuclear volumes remain similar to control (data not shown). To assess the effect of *Arp3* depletion on EC differentiation, we electroporated with sh*Arp3* and mb-cherry at P1 the lateral ventricles of Centrin 2-GFP expressing mice, and quantified the extent of differentiation at P5 and at P15. Already at P5, we observed that fewer progenitors had differentiated as compared to controls (Figure 2D, E). We should point out that some of the differentiated cells observed at P5 might have already been engaged in the process of differentiation at the time of electroporation as differentiation is asynchronous. This is consistent with the fact that at P15, we observed a much stronger defect in EC differentiation in cells electroporated with the sh*Arp3* compared to the control (Figure 2F, G). All together, these data suggest that branched F-actin is involved in the deformation / localization of the nuclei to promote EC differentiation.

**Figure 2:**
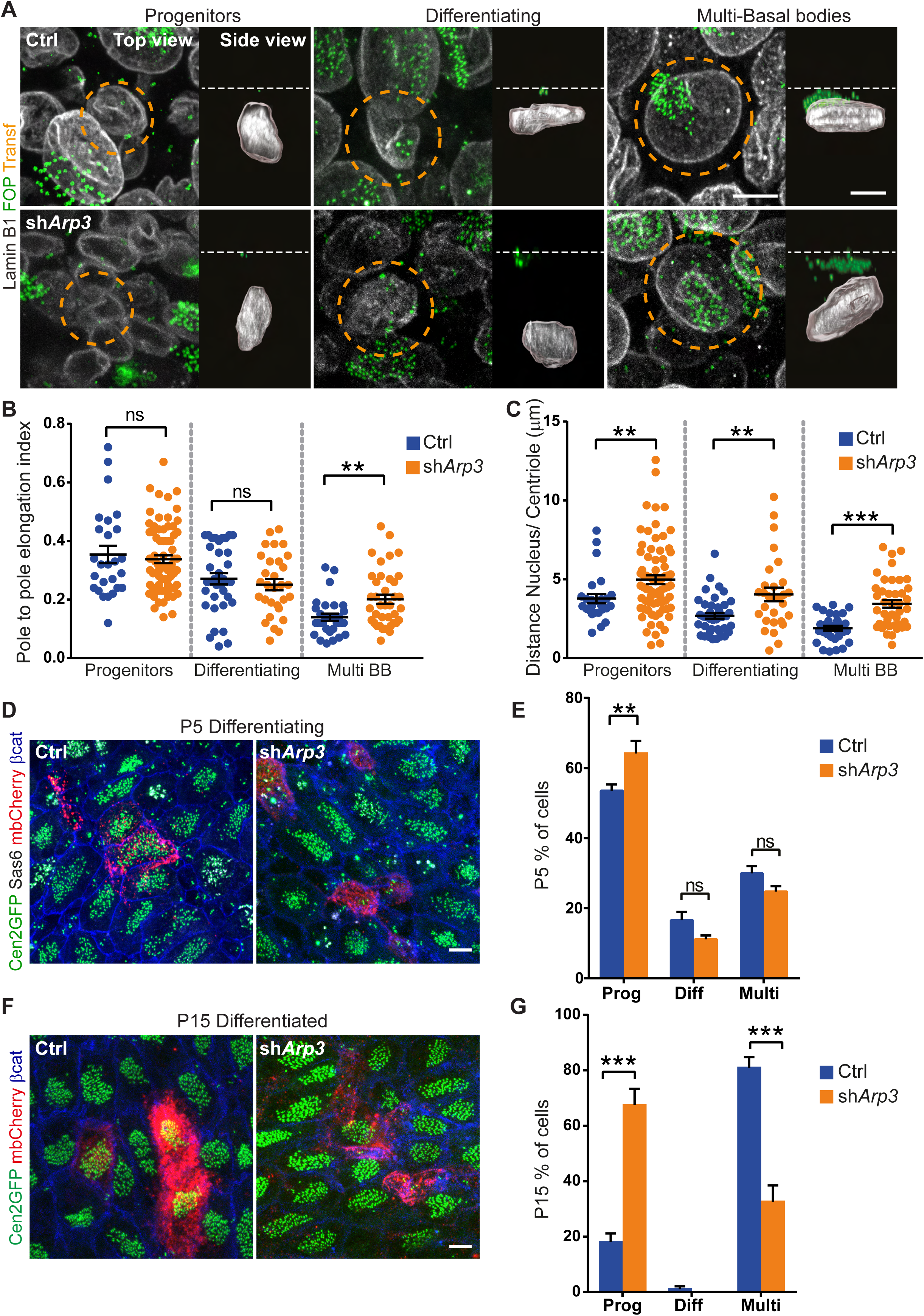
Decreasing ARP3 impairs nuclear deformation and ependymal differentiation. Wild-type **(A-C)** or Centrin2-GFP **(D, G)** mice transfected at P1 with mb-cherry as a reporter and either a control plasmid (Ctrl) or an shRNA against *Arp3* (sh*Arp3*). **A)** Staining of Lamin-B1 (nuclear lamina, Gray) and FOP (centrioles, Green) during ependymal cell differentiation (P5) on whole mount lateral wall. Transfected cells, either control (top row) or sh*Arp3* (bottom row), are circled with orange dashed lines. Top view is a maximum projection of z-stacks. Side view is an Imaris 3D projection with segmented nuclei. The white dashed lines represent the plasma membrane. **(B, C)** Quantification of nuclear shapes during the different steps of ependymal differentiation on Imaris segmented nuclei as in **A**. **B)** Pole to pole elongation index. **C)** Distance between the center of the nucleus and the highest centriole. Error bars represent the sem of n=26/ 73 progenitors, 40/ 29 differentiating and 31/ 39 multi-basal bodies (Multi BB) transfected cells (Ctrl/ sh*Arp3*) from four mice; p-values from Mann-Whitney test; *** P≤0.001; ** P≤0.01; ns, P>0.05. **D)** Staining of Centrin2-GFP (Cen2GFP, centrioles, Green), Sas6 (pro-centrioles, Gray), mb-cherry (reporter, Red) and beta-catenin (junction, Blue) during ependymal cell differentiation (P5) on whole mount lateral wall. **E)** Quantification of the percentage of transfected cells as in **D** at each stage of differentiation at P5 (Prog=progenitors, Diff=Differentiating, Multi=multi-basal bodies). Error bars represent the sem of 5 experiments n=174 (Ctrl) and 183 (sh*Arp3*) transfected cells; p-values from two-way ANOVA followed by Sidak’s multiple comparisons test; ** P≤0.01. **F)** Staining of Centrin2-GFP (Cen2GFP, centrioles, Green), mb-cherry (reporter, Red) and beta-catenin (junction, Blue) in differentiated cells (P15) on whole mount lateral wall. **G)** Quantification of the percentage of transfected cells as in **F** remaining as progenitors (Prog), still differentiating (Diff) or differentiated in ependymal cells (Multi), at P15. Error bars represent the sem of 4 or 3 experiments n=83 (Ctrl) and 93 (sh*Arp3*) transfected cells, respectively; p-values from two-way ANOVA followed by Sidak’s multiple comparisons test; *** P≤0.001. Scale bar = 5 μm.

If this were the case, we would expect that performing the inverse experiment, i.e. increasing actin polymerization, would increase nuclear deformation and EC differentiation. To test this hypothesis, we examined EC differentiation in an *Arpin* conditional KO mouse^49^; the *Arpin* gene encodes for a negative regulator of the Arp2/3 complex so the knockout has enhanced branched actin polymerization^50^. We electroporated at P1 the lateral ventricle of *Arpin* conditional KO mice with the Cre recombinase and the mb-cherry marker. We first confirmed that removing *Arpin* increased F-actin accumulation in differentiating ECs compared to control electroporation and relative to untransfected surrounding progenitors (Figure S3A, B). We also stained the electroporated lateral walls for Lamin-B1 and observed that increasing F-actin polymerization triggered nuclei deformation and migration toward the apical surface at earlier stages than control cells (Figure 3A-C). Finally, we assessed whether increasing F-actin perturbed EC differentiation by staining for centriole and deuterosome (the platform around which procentrioles are assembled) markers. We observed that removing *Arpin* at differentiation onset led to an increase in differentiated cells (Figure 3D, E) at P5. Similarly, at P15 we observed significantly more multiciliated ECs in the animals electroporated with Cre as compared to the control (Figure 3F, G). These results suggest that an increase in Arp2/3 activity led to premature differentiation of progenitors into ECs.

**Figure 3:**
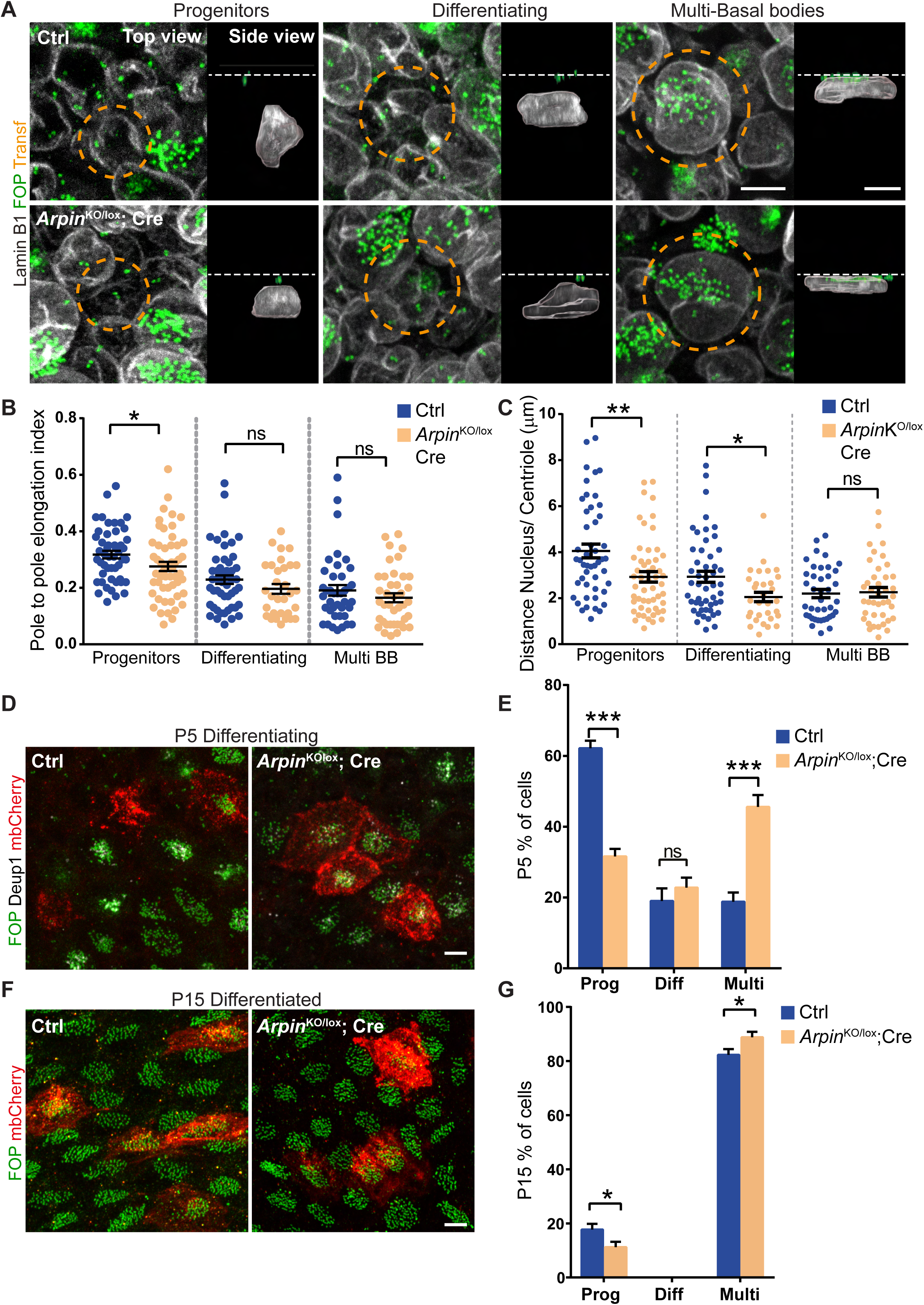
Increasing Arp2/3 complex activity facilitates nuclear deformation and ependymal differentiation. **(A-G)** Arpin^lox/lox^ transfected at P1 with mb-cherry as a reporter and a control plasmid (Ctrl) or Arpin^KO/lox^ mice transfected at P1 with mb-cherry as a reporter and a plasmid coding for the Cre recombinase (*Arpin*^KO/lox^; Cre). **A)** Staining of Lamin-B1 (nuclear lamina, Gray) and FOP (centrioles, Green) during ependymal cell differentiation (P5), on whole mount lateral wall. Transfected cells, either Ctrl (top row) or Arpin^KO/lox^; Cre (bottom row) are circled with orange dashed lines. Top view is a maximum projection of z-stacks. Side view is an Imaris 3D projection with segmented nuclei. The white dashed lines represent the plasma membrane. **(B, C)** Quantification of nuclear shapes during the different steps of ependymal differentiation on Imaris segmented nuclei of cells as in **A**. **B)** Pole to pole elongation index. **C)** Distance between the center of the nucleus and the highest centriole. Error bars represent the sem of n=48/ 53 progenitors, 50/ 31 differentiating and 40/ 40 multi-basal bodies (Multi BB) cells (Ctrl/ *Arpin*^KO/lox^; Cre) from 7 mice in Ctrl and 8 mice in *Arpin*^KO/lox^; Cre; p-values from Mann-Whitney test; ** P≤0.01; * p<0.05; ns, P>0.05. **D)** Staining of FOP (centrioles, Green), Deup1 (deuterosomes, Gray) and mb-cherry (reporter, Red) during ependymal cell differentiation (P5) on whole mount lateral wall. **E)** Quantification of the percentage of transfected cells as in **D** at each stages of differentiation at P5. Error bars represent the sem of 5 (Ctrl) or 4 (*Arpin*^KO/lox^; Cre) experiments, n=178 (Ctrl) and 145 (*Arpin*^KO/lox^; Cre) transfected cells; Prog=progenitors, Diff=Differentiating and Multi=Multi-basal body cells. p-values from two-way ANOVA followed by Sidak’s multiple comparisons test; *** P≤0.001; ns, P>0.05. **F)** Staining of FOP (centrioles, Green) and mb-cherry (reporter, Red) in differentiated cells (P15) on whole mount lateral wall. **G)** Quantification of the percentage of transfected cells as in **F** remaining as progenitors (Prog), still differentiating (Diff) or differentiated in ependymal cells (Multi) at P15. Error bars represent the sem of 6 (Ctrl) and 5 (*Arpin*^KO/lox^; Cre) experiments, n=314 (Ctrl) and 151 (*Arpin*^KO/lox^;Cre) transfected cells; p-values from two-way ANOVA followed by Sidak’s multiple comparisons test; * p=0.0416. Scale bar = 5 μm.

### Severing the link between the nucleus and the cytoskeleton perturbs nuclear deformation and ependymal differentiation

F-actin has many functions in cells so decreasing or increasing actin amounts could have off-target effects. To address this issue, we left the actin intact, but perturbed its link to the nucleus by expressing in the cells, at the onset of differentiation (P1), a dominant negative construct composed of the KASH domain from NESPRIN1^51^. This domain displaces the endogenous NESPRINS by interacting with the SUN proteins, but cannot interact with the cytoskeleton. We first assessed whether perturbation of the LINC complex changed F-actin organization during ependymal differentiation. We stained lateral ventricles electroporated at P1 with the KASH domain or a control plasmid and mb-cherry with phalloidin at P5. We did not observe changes in the F-actin organization compared to the control electroporated cells (Figure S3A, B) and relative to untransfected surrounding progenitor cells. However, considering the complexity of actin staining in differentiating cells, we cannot rule out some local actin rearrangement, especially as there are reports in other systems of crosstalk between the LINC complex and the actin cytoskeleton^52,53^. We also stained for Lamin-B1 and observed that the nuclei of differentiating cells (P5) expressing the KASH flattened less than control nuclei and remained in the depth of the tissue (Figure 4A-C). Thus, disrupting the LINC complex was sufficient to perturb nuclear morphology without drastically affecting F-actin polymerization. We then assessed how the cells differentiate in the absence of functional LINC complex. At P5, when the cells are differentiating, expressing the KASH domain already slightly impaired differentiation (Figure 4D, E), and this effect was even more pronounced at P15 (Figure 4F, G). All together, these experiments show that severing the link between the nucleus and the cytoskeleton is sufficient to perturb nuclear morphology and EC differentiation without affecting drastically actin organization.

**Figure 4:**
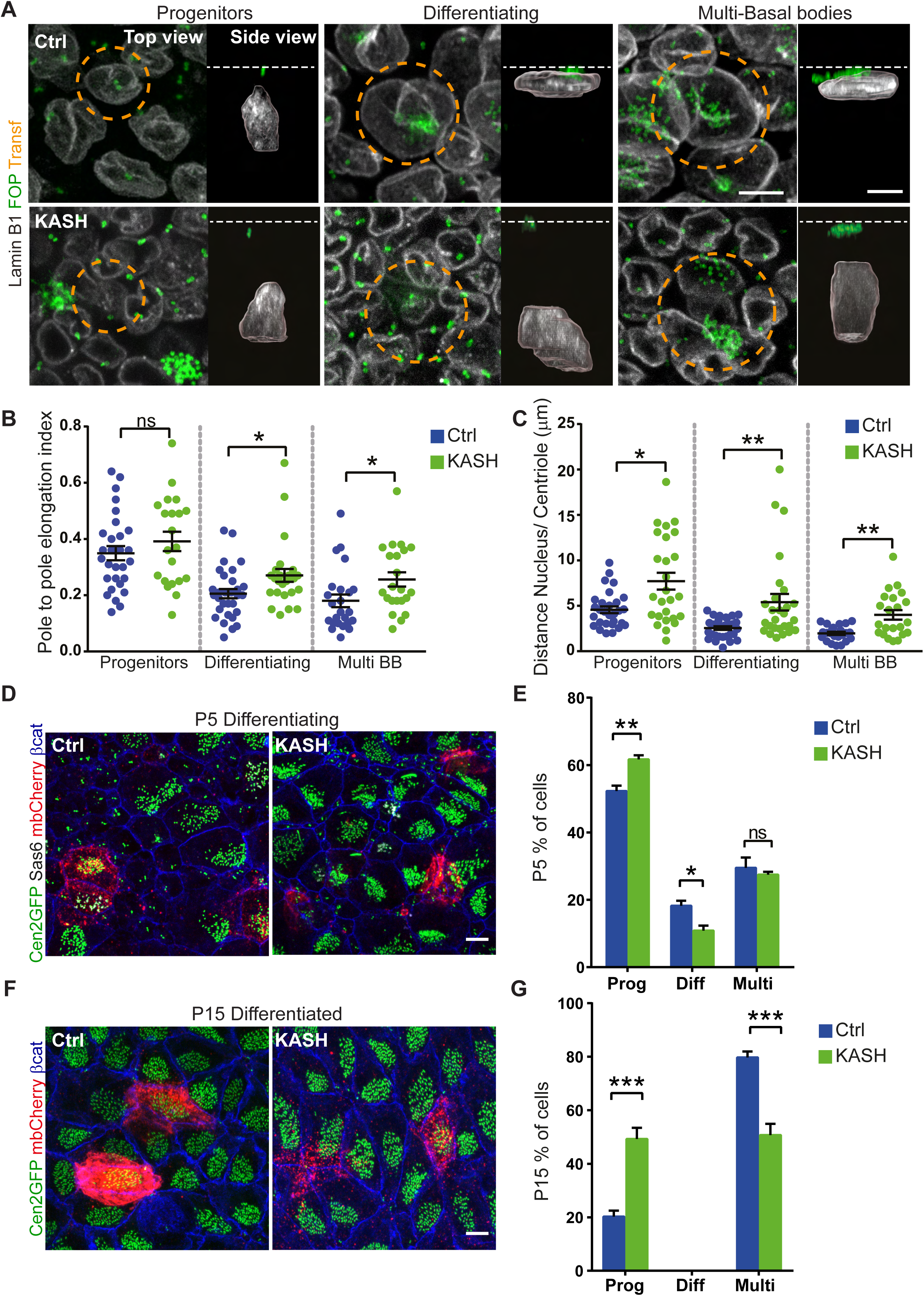
Severing the link between the nucleus and the cytoskeleton impairs nuclear deformation and ependymal differentiation. Wild-type **(A-C)** or Centrin2-GFP **(D-G)** mice transfected at P1 with mb-cherry as a reporter and either a control plasmid (Ctrl) or the KASH-domain (dominant negative of the LINC complex, KASH). **A)** Staining of Lamin-B1 (nuclear lamina, Gray) and FOP (centrioles, Green) during ependymal cell differentiation (P5) on whole mount lateral wall. Transfected cells, either Ctrl (top row) or KASH (bottom row), are circled with orange dashed lines. Top view is a maximum projection of z-stacks. Side view is an Imaris 3D projection with segmented nuclei. The white dashed lines represent the plasma membrane. **(B, C)** Quantification of the nuclear shape during the different steps of ependymal differentiation on Imaris segmented nuclei as in **A**. **B)** Pole to pole elongation index. **C)** Distance between the center of the nucleus and the highest centriole. Error bars represent the sem of n=29/ 21 progenitors, 30/ 26 differentiating and 23/ 22 multi-basal bodies (Multi BB) transfected cells (Ctrl/ KASH) from three mice; p-values from Mann-Whitney test * P<0.05; ns P>0.05; ** P≤0.01. **D)** Staining of Centrin2-GFP (Cen2GFP, centrioles, Green), Sas6 (procentrioles, Gray), mb-cherry (reporter, Red) and beta-catenin (junction, Blue) during ependymal cell differentiation (P5) on whole mount lateral wall. **E)** Quantification of the percentage of transfected cells as in **D** at each stages of differentiation at P5. Error bars represent the sem of 5 experiments; n=131 (Ctrl) and 189 (KASH) transfected cells; Prog=Progenitor, Diff=Differentiating, Multi=multi-basal body cells. p-values from two-way ANOVA followed by Sidak’s multiple comparisons test; ** P≤0.01; * P=0.035; ns P>0.05. **F)** Staining of Centrin2-GFP (Cen2GFP, centrioles, Green), mb-cherry (reporter, Red) and beta-catenin (junction, Blue) in differentiated cells (P15) on whole mount lateral wall. **G)** Quantification of the percentage of transfected cells as in **F** remaining as progenitors (Prog), still differentiating (Diff) or differentiated (Multi) at P15. Error bars represent the sem of 6 (Ctrl) or 3 (KASH) experiments, n=148 (Ctrl) and 86 (KASH) transfected cells; p-values from two-way ANOVA followed by Sidak’s multiple comparisons test; *** P≤0.001. Scale bar = 5 μm.

To extend this observation and identify which LINC complex components were involved, we stained the differentiating tissue with antibodies directed against LINC complex proteins. Among them, we saw that NESPRIN1 and 2 were expressed during differentiation (Figure 5A). We observed that NESPRIN1 accumulated at an early stage of differentiation at the centrosome/growing centrioles, and was enriched at the nuclear envelope at late stages of differentiation, while NESPRIN2 was recruited to the nuclear envelope very early in the differentiation process. We quantified this recruitment in an *in vitro* culture of differentiating progenitor cells^54-56^, as the background was less pronounced than *in vivo* (Figure S4A-D). This confirmed the appearance of NESPRIN2 at the nuclear envelope earlier than NESPRIN1 and the accumulation of NESPRIN1 at the centrosome during the process of differentiation. Intriguingly, we observed an increase in actin levels after NESPRIN1 depletion, using a validated shRNA sequence against Syne1^57^ (Figure S4E-G), in differentiating (Figure S3A, B) and progenitor cells (not shown). As it happened before NESPRIN1 was recruited to the nuclear envelope, it might dependent on the non-nuclear NESPRIN1. In accordance with the actin levels, NESPRIN1 depletion led to a premature flattening and re-localization of the nucleus (Figure S5A-C) and accelerated ependymal differentiation (Figure S5D-G). On the other hand, depleting NESPRIN2, using a validated shRNA sequence against Syne2^57^ (Figure S4H, I), had a similar effect as expressing the KASH domain: no drastic effect on actin organization (Figure S3A, B), and both nuclear deformation/re-localization (Figure 5B-D) and ependymal differentiation (Figure 5E-F) impairment. All together, these experiments show that severing the NESPRIN2/cytoskeleton interaction is sufficient to perturb the nuclear morphology and EC differentiation without drastically affecting actin organization.

**Figure 5:**
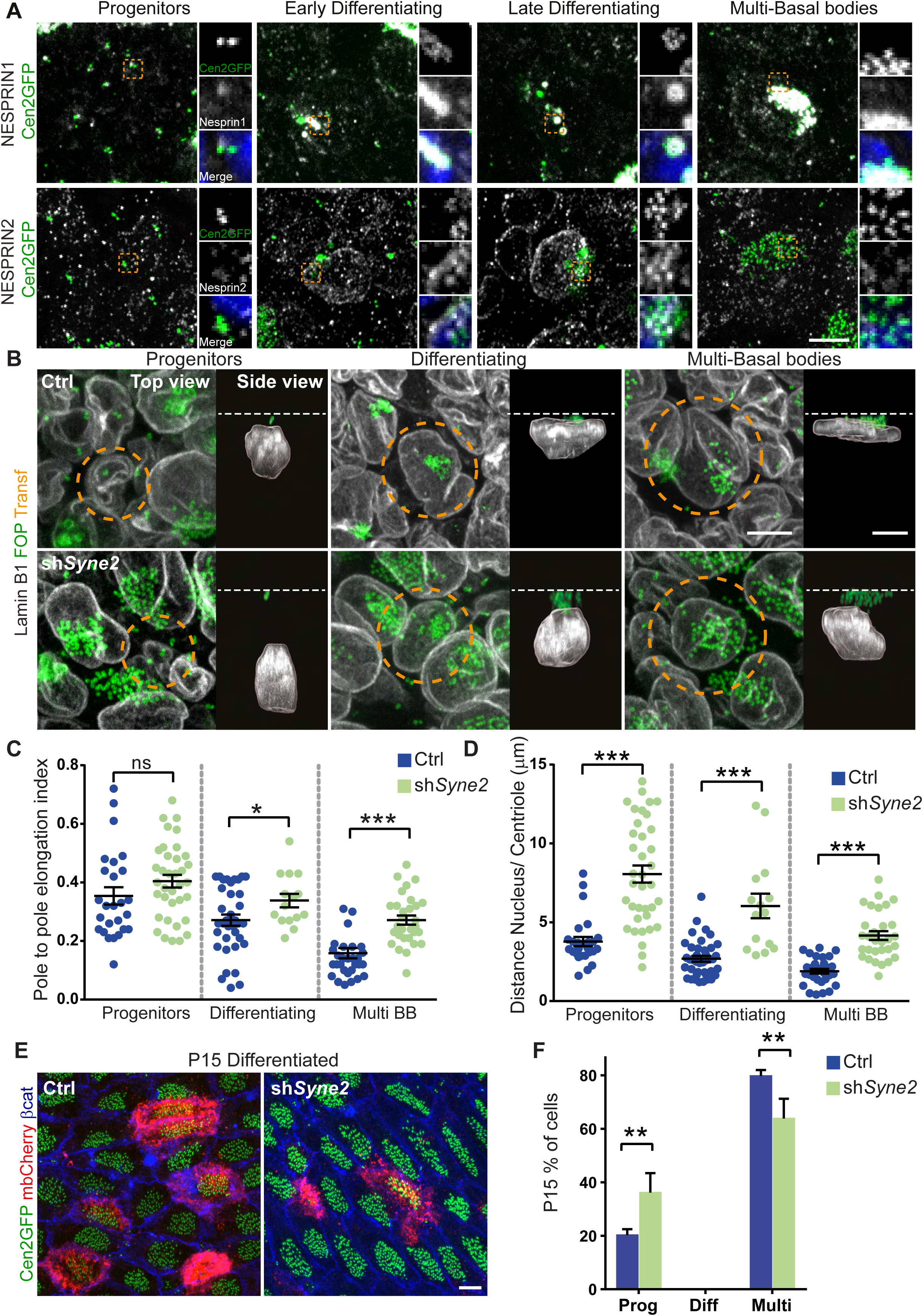
NESPRIN2 localizes earlier during differentiation at the nuclear envelope than NESPRIN1 and promotes nuclear deformation and ependymal differentiation. **A)** Staining of NESPRIN1 (Gray, top panel) or NESPRIN2 (Gray, bottom panel) and Centrin2-GFP (Cen2GFP, Centrioles, Green) during ependymal differentiation (P5) on whole mount lateral wall. Small insets are 5x single channel zoom images corresponding to the orange dashed square. Blue is Hoechst staining. Wild-type **(B-D)** or Centrin2-GFP **(E-F)** mice transfected at P1 with mb-cherry as a reporter and either a control plasmid (Ctrl) or the shRNA against *Syne2* (to deplete NESPRIN2). **B)** Staining of Lamin-B1 (nuclear lamina, Gray) and FOP (centrioles, Green) during ependymal cell differentiation (P5) on whole mount lateral wall. Transfected cells, either Ctrl (top row) or sh*Syne2* (bottom row), are circled with orange dashed lines. Top view is a maximum projection of z-stacks. Side view is an Imaris 3D projection with segmented nuclei. The white dashed lines represent the plasma membrane. **(C, D)** Quantification of the nuclear shape during the different steps of ependymal differentiation on Imaris segmented nuclei as in **B**. **C)** Pole to pole elongation index. **D)** Distance between the center of the nucleus and the highest centriole. Error bars represent the sem of n=26/ 37 progenitors, 40/ 15 differentiating and 31/ 30 multi-basal bodies (Multi BB) transfected cells (Ctrl/ KASH) from three mice; p-values from Mann-Whitney test * P<0.05; ns P>0.05; ** P≤0.01. Control values are the same than in Figure 2 as the experiments were done at the same time. **E)** Staining of Centrin2-GFP (Cen2GFP, centrioles, Green), mb-cherry (reporter, Red) and beta-catenin (junction, Blue) in differentiated cells (P15) on whole mount lateral wall. **F)** Quantification of the percentage of transfected cells as in **E** remaining as progenitors (Prog), still differentiating (Diff) or differentiated (Multi) at P15. Error bars represent the sem of 6 (Ctrl) or 3 (sh*Syne2*) experiments, n=148 (Ctrl) and 84 (sh*Syne2*) transfected cells; The control values are the same than the one in Figure 4 as the experiments were performed at the same time. p-values from two-way ANOVA followed by Sidak’s multiple comparisons test; *** P≤0.001. Scale bar = 5 μm.

### Confining the cell and its nucleus induces EC differentiation

To more directly assess whether nuclear deformation triggered ependymal differentiation, we applied an artificial mechanical load on the nucleus. As it is challenging to perform mechanical loading on cells *in vivo*, we used our cell culture model of ependymal differentiation. We first confirmed that the nuclei deformed during *in vitro* differentiation. To do so, we stained Centrin2-GFP cells 5 days (D5) after serum withdrawal (differentiation onset, D0) for Lamin-B1 (lamina) and Sas6. We then analyzed the nuclear shape at different stages of differentiation as determined by the pattern of Centrin2-GFP/ Sas6. We observed that *in vitro* the nuclei increased in size (1.3x) and flattened (1.4x) during ependymal differentiation (Figure S6A-C), as *in vivo*. Additionally we observed that H3K27me3 increased during *in vitro* differentiation while H3K9me2 decreased (Figure S6D-F). We also performed shRNA against *Arp3* and *Syne2 in vitro* and observed that their depletion led to an impairment of EC differentiation (Figure 6A, B). Interestingly, depleting NESPRIN2 led to an increase of H3K9me2 levels in the progenitor cells, indicating that forces applied on the nucleus are dependent on NESPRIN2^43^ (Figure S6G, H). Thus, *in vitro,* like *in vivo,* the link between the nucleus via NESPRIN2 and the actin cytoskeleton is important for EC differentiation.

**Figure 6:**
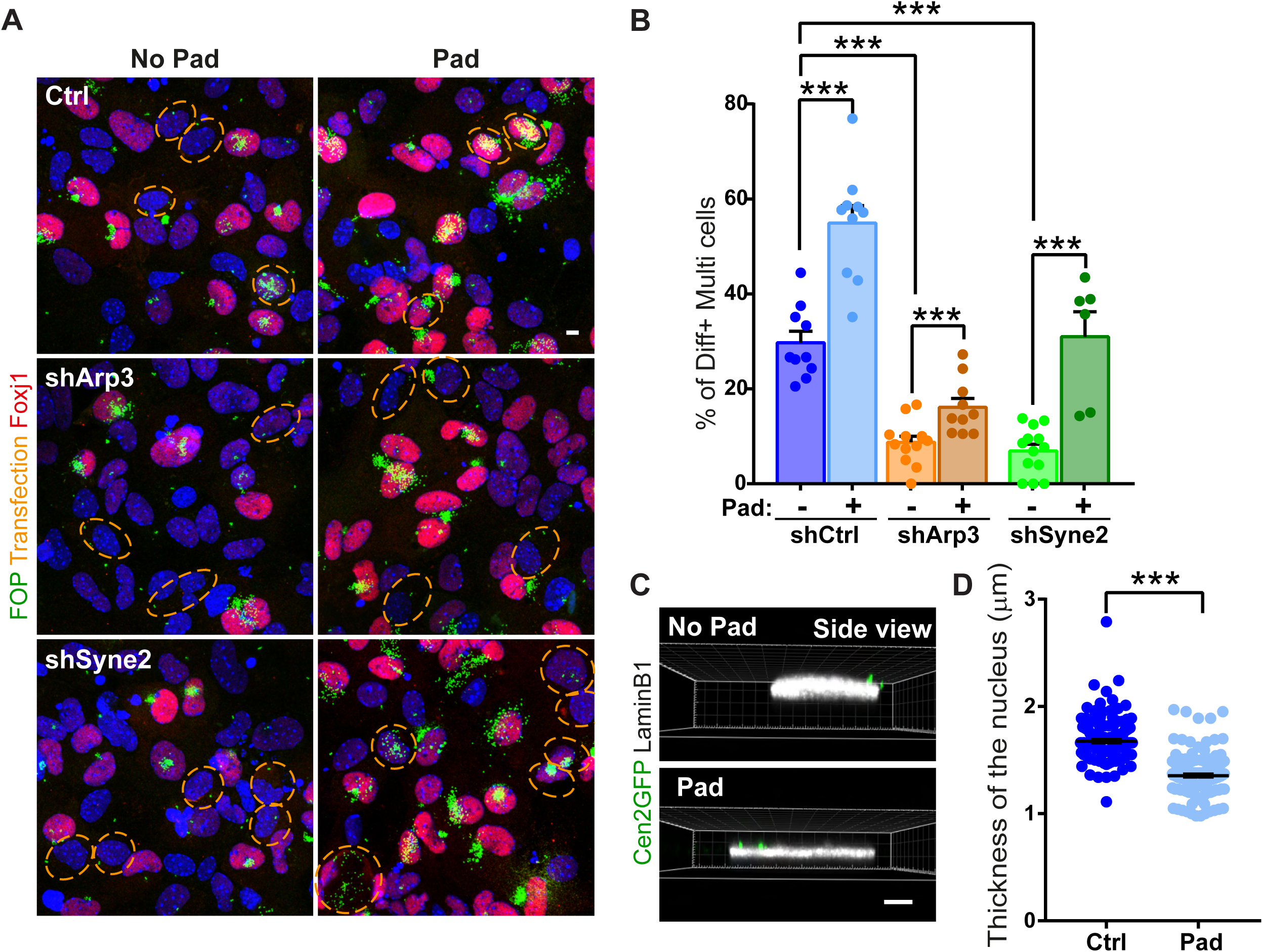
Confining the progenitor cells *in vitro* induces ependymal differentiation and rescues the absence of NESPRIN2. **(A, B)** Progenitor cells *in vitro* were transfected with shRNA either control (top panel) or against *Arp3* (middle panel) or against *Syne2* (bottom panel) at differentiation onset (D0= serum removal). They were then confined (Pad) or not (No Pad) under an agarose pad for 6 days. **A)** Staining of transfected control or confined cells 6 days after differentiation onset with FOP (centriole, Green), Foxj1 (transcription factor specific of multiciliated differentiating and differentiated cells, Red) and Hoechst (DNA, Blue). Transfected cells were circled with orange dashed line. The same image with the tomato reporter can be found on Figure S6I. **B)** Quantification of the percentage of differentiating (Diff) + differentiated (Multi) transfected cells 6 days after the confinement and shRNA expression. Error bars represent the sem of at least 6 coverslips (over 3 independent experiments) and over 115 transfected cells. p-values from Mann-Whitney test. *** P≤0.001. **(C, D)** Progenitor cells *in vitro* were confined (Pad) or not (No Pad) under an agarose pad from differentiation onset (D0) for 6 days. **C)** 3D Imaris view from the side of the cells. **D)** Quantification of the nucleus thickness of progenitor cells with Imaris software on image as in **C**. Error bars represent the sem of 156/ 165 nuclei measured on three independent experiments. p-value from Mann-Whitney test, *** P≤0.001. Scale bar = 5 μm.

To compress the nucleus, we overlaid cells with an agarose pad at differentiation onset (D0), as previously used for other cell types^29,58^. After 6 days, we measured the height of the nuclei on confined and control cells stained with Lamin-B1 and observed a significant flattening (Figure 6C, D). We also counted the number of differentiating/differentiated cells and observed an increase in their number following compression (Figure 6A, B). Thus, mechanical load on the cell is sufficient to deform the cell nucleus and induce ependymal differentiation. To evaluate whether mechanical loading compensated for deficiencies in the LINC complex and actin, we compressed cells depleted for *Arp3* or *Syne2* (Figures 6A, B and S6I). In both cases, we observed increased differentiation in depleted cells under compression as compared to un-compressed cells. Taking the *in vitro* and *in vivo* experiments together, we conclude that nuclear deformation and EC differentiation go hand-in-hand, with nuclear deformation seemingly triggering EC differentiation.

### S-phase transition factors are downstream of nuclear deformations during EC differentiation

This result begs the question as to what is transducing the mechanical deformation of the nucleus to the transcription factors that are known to control the differentiation of multiciliated cells. There is extensive evidence in other systems that the transcription factor Yes-Associated Protein (YAP) is involved in mechanotransduction^27,59^. We therefore stained lateral ventricles at P5 with YAP/ TAZ antibody, and showed an increase of the YAP/ TAZ levels in the cytoplasm as well as a translocation of these proteins into the nucleus during ependymal differentiation (Figure S7A-C). However mice mutant for YAP (YAP^lox/lox^; Nestin:Cre depleted from E11.5) still displayed ependymal markers (cilia and centrioles) at both P5 and P10, like control litter mates, suggesting that YAP is not required for triggering ependymal differentiation on the lateral ventricles (Figure S7D).

In searching for an alternative mechanotransduction pathway, we noticed that in other systems, mechanical strain or cell confinement triggers cell cycle events and notably the transition between G1 and S phases^29,30,33^. In this cycling cells, CDK2 is known to phosphorylate retinoblastoma protein (RB1), triggering the release of E2F transcription factors from their interaction with RB1 and allowing them to activate S-phase specific genes to transition out of G1^60^. In ependymal cells, we know from previous work that CDK2 is also required for the early steps of multiciliated differentiation^11,61^ and that RB1 is hyperphosphorylated in differentiating EC^61^. Moreover, E2F4 and 5, along with the master regulators GEMC1 and MCIDAS, are required to activate transcription of specific genes for multiciliated cell differentiation^9^. Finally, RBL2 in the airway inhibits differentiation through Mcidas^62^. In keeping with this, expression of a non-phosphorylable version of RB1^63^ (hRB1ΔCDK) gave reduced differentiation as compared to control cells (Figure 7A, B). This is unlikely to be due to a change in nuclear deformation as H3K27me3 still increased during the differentiation (Figure S8A, B). Furthermore, we observed that MCIDAS overexpression did not induce rapid ependymal differentiation in cells expressing non-phosphorylable RB1 (Figures 7C, D and S8C), suggesting that RB1 phosphorylation/ release of E2F is important for MCIDAS activation during differentiation. All together, these results suggest that the phosphorylation of RB1 is required for ependymal differentiation through MCIDAS activation.

**Figure 7:**
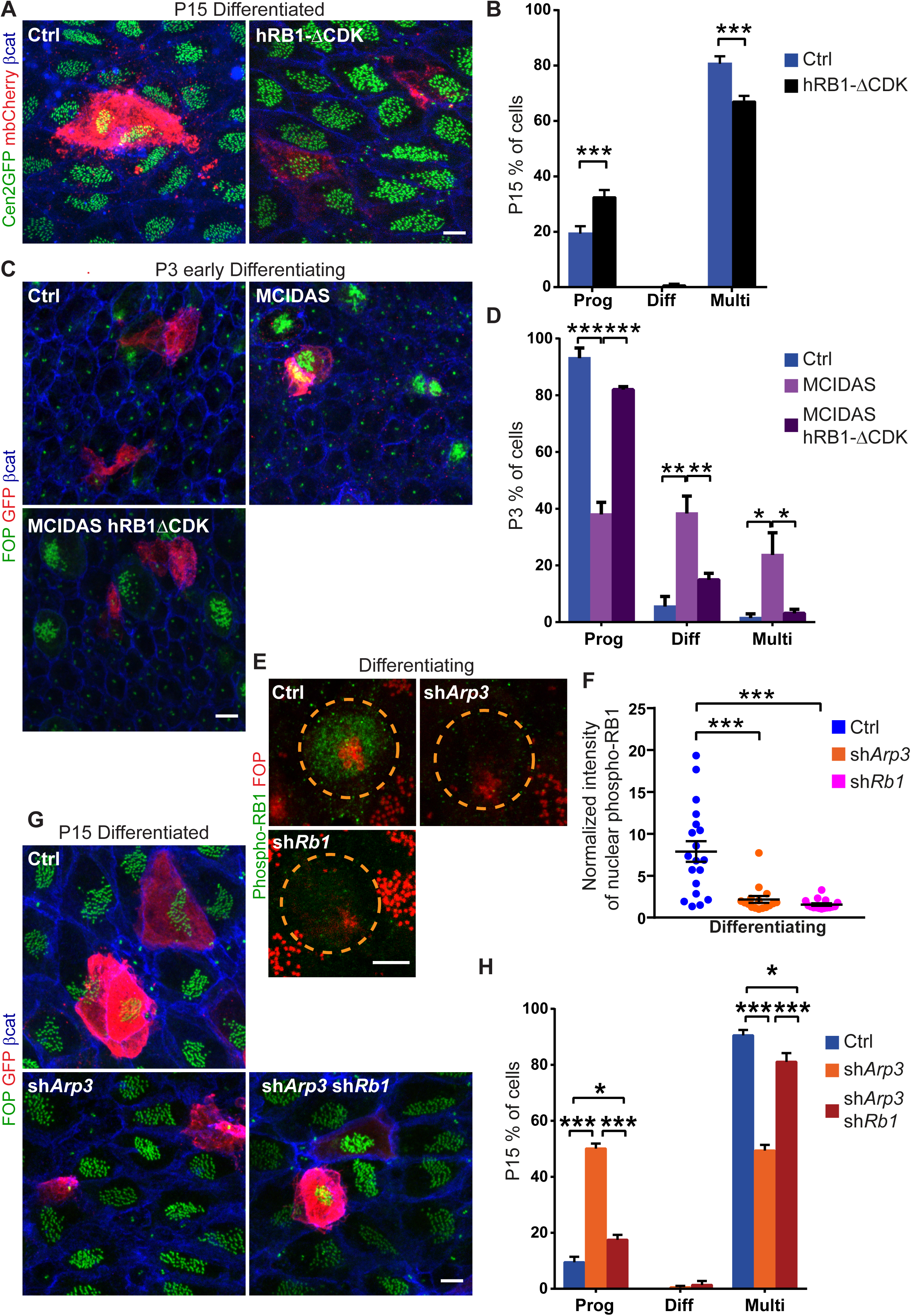
RB1 is involved in ependymal differentiation following nuclear deformation. **A)** Staining of Centrin2-GFP (Cen2GFP, centrioles, Green), mb-cherry (reporter, Red) and beta-catenin (junctions, Blue) in differentiated cells (P15) on whole mount lateral wall of Centrin2-GFP mice transfected at P1 with mb-cherry and either a control plasmid (Ctrl) or a plasmid coding for an unphosphorylable RB1 (hRB1-ΔCDK). **B)** Quantification of the percentage of transfected cells as in **A** remaining as progenitors (Prog), still differentiating (Diff) or differentiated (Multi) at P15. Error bars represent the sem of 4 experiments, n=146 (Ctrl) and 148 (hRB1-ΔCDK) transfected cells; p-values from two-way ANOVA followed by Sidak’s multiple comparisons test; *** P≤0.001. **C)** Staining of FOP (centrioles, Green), GFP (reporter, Red) and beta-catenin (junctions, Blue) in early differentiating cells (P3) on whole mount lateral wall of wild-type mice transfected at P1 with control plasmids (Ctrl= shNeg-IRES-GFP+ shNeg) or MCIDAS (shNeg-IRES-GFP+ MCIDAS) or both MCIDAS and unphosphorylatable RB1 (hRB1-ΔCDK –IRES-GFP+ MCIDAS). **D)** Quantification of the percentage of transfected cells at P3 as in **C** in the different stages of differentiation: Progenitors (Prog), differentiating (Diff) or differentiated (Multi). Error bars represent the sem of 3 (Ctrl) or 8 (MCIDAS) or 5 (MCIDAS + hRB1-ΔCDK) experiments, n=75 (Ctrl), 198 (MCIDAS) and 123 (MCIDAS + hRB1-ΔCDK) transfected cells; p-values from two-way ANOVA followed by Sidak’s multiple comparisons test; *** P≤0.0001; ** p≤0.01; * p≤0.05. **E)** Staining of phospho-RB1 (Green) and FOP (centrioles, Red) in differentiating cells transfected with mb-cherry (orange dashed circle) and either a control plasmid (Ctrl, top row), an shRNA against *Arp3* (sh*Arp3*) or an shRNA against *Rb1* (sh*Rb1*). **F)** Quantification of the fluorescence intensity of nuclear phospho-RB1 of transfected cells as in **D** normalized to the mean intensity of nuclear phospho-RB1 in three surrounding untransfected progenitors. Error bars represent the sem of n=19/ 16/ 16 (Ctrl/ sh*Arp3*/ sh*Rb1*) transfected cells. p-values from Mann-Whitney test. *** p≤0.001, * p=0.0125. **G)** Staining of FOP (centrioles, Green), GFP (reporter, Red) and beta-catenin (junction, Blue) in differentiated cells (P15) on whole mount lateral wall of wild-type mice transfected with a control plasmid (Ctrl= shNeg-IRES-GFP+ shNeg) or sh*Arp3 (shArp3*-IRES-GFP+ shNeg) or both sh*Arp3* and sh*Rb1 (shArp3*-IRES-GFP*+ shRb1)*. IRES containing constructs were used as reporter, less concentrated than the other plasmids (potentially explaining the decreased effect of sh*Arp3* compared to Figure 2). **H)** Quantification of the percentage of transfected cells as in **A**, remaining as progenitors (Prog), still differentiating (Diff) or differentiated in ependymal cells (Multi), at P15. Error bars represent the sem of 6 (Ctrl) or 6 (sh*Arp3*) or 3 (sh*Arp3* + sh*Rb1*) experiments, n=109 (Ctrl), 155 (sh*Arp3*) and 68 (sh*Arp3*+ sh*Rb1*) transfected cells; p-values from two-way ANOVA followed by Sidak’s multiple comparisons test; *** p≤0.001; for Prog Ctrl vs sh*Arp3*+ sh*Rb1* *p=0.0483; for Multi Ctrl vs sh*Arp3*+ sh*Rb1* *p=0.0151. Scale bar = 5 μm.

A possibility, therefore, is that nuclear deformation during differentiation leads to the phosphorylation/ degradation of RB1 releasing E2F to activate MCIDAS. To test this hypothesis, we depleted *Arp3* in cells of the lateral wall (from differentiation onset) a treatment that gave impaired nuclear deformation and ependymal differentiation (Figure 2). We observed that under these conditions phospho-RB1 failed to accumulate in the nucleus of differentiating cells as compared to control (Figure 7E, F). Conversely, we rescued differentiation in *Arp3* depletion conditions by co-depleting *Rb1* (Figure 7G, H) with previously validated shRNA^64^ (Figure 7E, F). Note that *Rb1* depletion had no effect on differentiation on its own (Figure S8D, E).

Putting all this data together, we conclude that RB1 phosphorylation is a likely player in the pathway connecting actin accumulation and nuclear deformation to changes in differentiation.

## DISCUSSION

Impairing nuclear deformation either by decreasing actin (sh*Arp3*, Figure 2) or by severing the link between the nucleus and the actin cytoskeleton (KASH expression or sh*Syne2* Figures 4, 5), impaired EC differentiation. Conversely, forcing nuclear deformation by increasing actin polymerization (*Arpin^KO/lox^*; Cre or shSyne1) (Figures 3, S3 and S5) or compressing the cells (Figure 6) accelerated differentiation. Our results thus show that mechanical constraints on the nucleus are important for EC differentiation in the mouse brain. In line with the presence of mechanical forces on the nucleus, Histone H3 methylation (Figures 1E-G and S6D-F), and DNA-damage (not shown^61^) are observed during differentiation, similarly to reports in other cell types submitted to mechanical stress^35,43^. Interestingly, compression of the nucleus rescued the effect of *Syne2* depletion on differentiation to the levels of control without compression while it only partly re-established differentiation in cells depleted for Arp3 (Figure 6). This was consistent with actin playing a role even when not linked through the LINC complex to the nucleus.

Independently of the Hippo pathway, it was shown that YAP entry in the nucleus is sensitive to forces applied on the nucleus through the opening of nuclear pores^27^. Interestingly, the YAP pathway is also involved in cell proliferation and differentiation in several tissues^59^. In the lung, depletion of YAP during cell specification prevents the formation of multiciliated cells at the expense of progenitor and goblet cells. However, at differentiation onset, expression of a nuclear YAP prevents lung multiciliated cell differentiation^65,66^. Similarly in the brain, YAP is involved in the specification/ integrity of ependymal progenitor cells in the aqueduct and to a lesser extend in the cortex^67^. Consequently, YAP absence leads to the blockage of the rostral aqueduct and non-communicating hydrocephalus. However, removing YAP does not prevent ependymal differentiation (Figure S7D).

In cycling cells, CDK2 is known to phosphorylate retinoblastoma protein (RB1), triggering the release of E2F transcription factors from their interaction with RB1 and allowing them to activate S-phase specific genes to transition out of G1^60^. Interestingly, mechanical strain or cell confinement also triggers cell cycle events and notably the transition between G1 and S phases^29,30,33^. Whether this depends on RB1 phosphorylation was not assessed. Multiciliated cell differentiation depends on the master regulators GEMC1 and MCIDAS that together with E2F4 or 5 activate transcription of specific genes^9^. It also depends on CDK2^11^ and RB1 is hyperphosphorylated^61^, even if the cells are post-mitotic. We show here that RB1 phosphorylation is important for ependymal differentiation in a pathway involving MCIDAS (Figure 7A-D) and propose that RB1 phosphorylation could provide a possible explanation for how nuclear deformation impacts transcription factor activity for differentiation. Indeed, removal of RB1 rescued ependymal differentiation in absence of nuclear deformation (sh*Arp3*, Figure 7G, H). Moreover, RB1 phosphorylation depends on nuclear deformation (Figure 7E, F). Thus, nuclear deformation may induce RB1 phosphorylation that will allow the release of E2F present in the cells and the activation of the multiciliated master regulators. Cells will remain in undifferentiated state until the nucleus is constrained by the actin cytoskeleton. This constraint will then induce the first steps of cell cycle-like activation that will trigger ependymal differentiation without inducing DNA replication^61^.

At this stage, we cannot rule out other mechanisms linking nuclear deformation to differentiation. It is for example unclear whether the changes in histone 3 methylation patterns (going from H3K9me2 to H3K27me3, Figures 1E-G and S6D-F), that depends on the LINC complex (Figure S6G, H), is necessary/ sufficient for the differentiation. Interestingly, RB1 interacts with HP1/ SUV39H1 that are essential for H3K9 methylation^68^. Thus, RB1 phosphorylation might influence H3K9 methylation. Similarly, nuclear pore or nuclear channel opening following nuclear deformation may play a role in the differentiation^69^, even if it does not imply YAP in our model (Figure S7).

The mechanical cues leading to actin accumulation that will act on the nucleus are not known. Neural stem cell progenitors are submitted to a variety of forces ranging from tissue growth (not accounted by division^7^), changes in morphology (going from elongated to more cuboidal during differentiation) and pressure or shear stress generated by the cerebrospinal fluid^12,70^. Interestingly, stretch activated channels, like PIEZO1 are involved in neural stem cell differentiation^71^ and may contribute to some aspect of multiciliated cell differentiation^72^.

## ACKNOWLEDGEMENTS

The data that support the findings of this study are available within the Article and Supplemental Files, or available from the authors upon request. All unique/stable reagents generated in this study are available from the lead contact. We are grateful to J. Gleeson for providing the Centrin2-GFP mice. We thank X. Morin for the CAAGS-mb-cherry and CAAGS-Cre plasmids, I. Caillé and S. Scotto for the shRNA backbones, A. Merdes for the KASH plasmid, S. Dowdy for the hRB1ΔCDK plasmid. G. Gundersen for the NESPRIN2 antibody, S. Taraviras for the Mcidas plasmid, NJ Quintyne for the p150-CC1 plasmid, M. Pende and E.N. Olson for the Yap^lox^ mice. We are grateful to A. Delecourt, E. Touzalin, G. Firmin and L. Le Moal for their help with the animal husbandry. We thanks M. Bornens for all his input over the year. We are grateful to all members of the Spassky laboratory.

The team was funded by the Agence Nationale de la Recherche Investissements d’Avenir (ANR-10-LABX-54MEMO LIFE, ANR-11-IDEX-0001-02 PSL* Research University, ANR-10-IDEX-0001-02 PSL, and ANR-10-LABX-31). The Spassky team is supported by the Inserm, the CNRS, the ENS, the ANR (ANR-20-CE45-0019, ANR-21-CE16-0016, and ANR-22-CE16-0011), the Fondation pour la Recherche Médicale (FRMEQU202103012767), the Fondation Pierre-Gilles de Gennes (FPGG03) and the European Research Council (ERC Consolidator grant 647466).

## AUTHOR CONTRIBUTIONS

Investigation and Formal Analysis: M.B., A.Mahuzier, S.K.A., A.Marty, A.S., M.K.D and N.D. Conceptualization: M.B., A.Mahuzier and N.D., Methodology and Resources: A.M.L.D., M.F. and A.B., Writing and Supervision: J.P., N.S., A.Y. and N.D., Funding Acquisition: A.Meunier, A.Y., J.P., N.S. and N.D.

## SUPPLEMENTAL INFORMATION

Document S1: Figure S1-S9

## MATERIALS AND METHODS

### Transgenic mice

All animal studies were performed in accordance with the guidelines of the European Community and French Ministry of Agriculture and were approved by the Ethic comity Charles Darwin (C2EA-05) and “Direction départementale de la protection des populations de Paris”, (Approval number Ce5/2012/107; APAFiS #9343). Both male and female animals were used in the study. Most of the mouse strains have already been described: wild-type are RjOrl:SWISS (Janvier labs), Cen2–GFP^38^ (B6D2-Tg(CAG-EGFP/CETN2)3-4Jgg/J, The Jackson Laboratory); B6D2-Arpin conditional knockout mice (Arpin^lox^) were generated at CIPHE (Centre d’Immunophénomique, Marseille, France) using CRISPR cas-9 technology^49^. C57BL/6-YAP conditional knockout (Yap^lox/lox^)^73^ were crossed with C57BL/6-Tg(Nestin-Cre)^74^; Yap^lox/+^.

### Plasmids

We cloned the shRNA sequences in the shRNA backbone vectors pcDNA-U6-shRNA and pTRIP-U6-shRNA-IRES-PGK-GFP-WPRE as previously described^75^. The control shRNA (shCtrl) was generated from AllStars Negative Control siRNA (Qiagen), which does not recognize mammalian RNAs. The reporter plasmid encoding membrane Cherry (mb-cherry) under the control of the CAGGS promoter^76^ was co-transfected with pcDNA-U6-shRNA and was a gift from Xavier Morin. All cells expressing the shRNA-GFP (pTRIP-U6-shRNA-IRES-PGK-GFP-WPRE) or cells expressing pCAGGS-mbCherry (mb-cherry), which were co-transfected with the untagged pcDNA-U6-shRNA, were considered to be depleted. shRNA sequences were validated in referenced paper and in this study: shArp3^48^: TTAGCTCTCTTCTACATCTGC (Figure S3A, B); shSyne2^57^: AGCCACAGAACTCCAAAAT (Figure S4H, I); shSyne1^57^: GCTCCTGCTGCTGCTTATT (Figure S4A, G); shRb1^64^: AGTTAGGACTGTTATGAAT (Figure 7D, E)).

The pCAGGS-Cre^77^ plasmid is gift from X. Morin. The KASH domain construct is a gift from A. Merdes^57^ and is tagged with GFP. The p150Glued-CC1 domain construct is a gift from NJ Quintyne^78^ and is tagged with dsRed. The pCMV HA hRB1 delta CDK was a gift from S. Dowdy (Addgene plasmid # 58906; http://n2t.net/addgene:58906; RRID:Addgene_58906)^63^. HA hRB1 delta CDK was subcloned at BamHI restriction site on pIRES2-EGFP vector (Clontech). The Mcidas-GFP construct was a gift from S. Taraviras^79^, in our hand the GFP was barely detectable. pCMV-tdTomato is from Clontech.

### Postnatal electroporation

Electroporation was performed on one-day-old pups as previously described^80^. shRNA in the U6 backbone (untagged) (2.5 μg) were co-transfected with the reporter plasmid mb-cherry or with plasmids encoding shRNA in the pTRIP backbone (IRES GFP) (1.5 µg). MCIDAS-GFP (2 μg) was electroporated either with pTRIP-shNeg or with HA hRB1 delta CDK-IRES2GFP (2 μg). Cherry-LmnA (1.5 μg) was electroporated either with pcDNA-U6-shNeg (2μg) or pcDNA-U6-shRb1 (2μg). All cells expressing the reporter were assumed to express both the reporter and the plasmid of interest.

### Immunostaining

Lateral walls of the lateral brain ventricles were dissected, fixed, and immunostained as previously described^54^. The same area (rostro-dorsal) of the lateral walls was used throughout the study. The tissue was treated with 0.1% triton in BRB80 (80 mM K-Pipes pH6.8; 1 mM MgCl2; 1 mM Na-EGTA) for 1 min and fixed in methanol at −20°C for 10 min, most of the time. For phalloidin stained tissues, they were detergent-treated and then fixed in 4% PFA in BRB80 for 8 min, for phospho-RB1, phosphor-Lamin-A and H3K37me3 staining or tissue electroporated with an IRES-GFP containing construct, the tissue were fixed in 4% PFA in BRB80 for 20 minutes. The following antibodies were used: rabbit or rat anti-dsRed (1:500, Clontech 632496 or 5F8 1:400, Chromotek 5f8-100); chicken anti-GFP (1:1600, Aves GFP-1020); mouse IgG2b anti-FOP (FGFR10P, 1:1000, Abnova H00011116-MO), mouse IgG1 anti-beta-catenin (1:1000, Millipore05-665), mouse IgG2b anti-Sas6 (1:700, Santa-Cruz Sc-81431), mouse IgG1 anti-LaminB1 (1:1000, Santa-Cruz Sc-374015), Rabbit anti-Deup1 (1:2000) ^81^, Rabbit anti-phospho S807/811-RB (1:1000, Cell Signaling 8516), mouse IgG1 anti-Foxj1 (2A5, 1:1000, Thermo-Fisher 14-9965-82), rabbit anti-Nesprin2 (1/100, gift from Greg Gundersen^82^), rat anti-tubulin YL1/2 (1:100, Sigma MAB1864), mouse IgG2a anti-Centrin (20H5 1:1000, Sigma 04-1624), mouse IgG1 anti-Nesprin-1 (1:50, Hybridoma bank MANES1A), mouse IgG2a anti-H3K9me2 (1:200, abcam ab1220), rabbit anti-H3K27me3 (1:100, Cell Signaling 9733S), rabbit anti-YAP/TAZ (1:50, D24E4 Cell Signaling 8418), mouse IgG1 polyglutamylation modification (GT335, 1:1000, Adipogen AG-20B-0020), and species-specific Alexa Fluor secondary antibodies (1:800, Invitrogen). Phalloidin-A488 or A568 were used at a dilution of 1:50 with primary and secondary antibodies.

### Microscopy

#### Optical imagery

Images (230-nm z-steps) of fixed samples were obtained with an inverted LSM 880 Airyscan (Zeiss) with a 63x 1.4NA oil-immersion objectives on confocal mode, except for the phalloidin staining where it was used in the airyscan mode (180nm z-steps), and Zen2 software. Z-stack projections are shown. Only the 7 most apical planes are shown for phalloidin staining. For P150-CC1 and KASH nuclei some intermediate Z were removed to allow visualisation of the transfected nuclei in the depth of the tissue. The KASH-GFP or the Cherry-LmnA around the nucleus are not visible on images representative of EC differentiation as the Z-stack including the apical surface does not include the whole nuclei deeper in the tissue.

### Drug treatments

Lateral wall explants were immersed for the indicated times at 37°C with 5% CO_2_ in DMEM-Glutamax (Invitrogen) supplemented with 10% FCS, 1% penicillin/streptomycin (PS) and cytochalasin-D (2μM, Sigma) to depolymerise actin, SMIFH2 (60μM, Sigma) to inhibit formin^83^, CK666 (100μM, Sigma) to inhibit the Arp2/3 complex, Blebbistatin (100μM, Sigma) or Nocodazole (10μM, Sigma) to depolymerize microtubules. The lateral ventricular wall of the opposite side of each mouse brain were used as control and immersed in the same media containing an amount of DMSO equivalent to that of the drug.

### Cell cultures

Differentiating ependymal cells were cultured as previously described^54^.

### Agarose pad

To compress the nuclei, we prepared 5mL of 2% ultra-pure agarose (FMC Bio-Products) in serum-free DMEM medium in a 60-mm Petri-dish. We made sure that the plates were on a flat surface before to pour the agarose. A 33-mm diameter agarose pad was cut out and deposited gently on cells grown on coverslips. A 32-mm glass coverslip surmounted of a 540 mg washer was carefully added above to maintain the cushion for compression^29^.

### *In vitro* transfection

We transfected 10ng of the plasmid of interest with 10ng of reporter (pCMV-tdTomato) on 0.2×10^6^ cells in suspension (before D0) with 0.04 μL of Jetprime (Polyplus) according to manufacturer protocol. The cells were seeded onto a drop of 20μL on a coverslip coated with poly-lysine. 500 μL of DMEM-Glutamax complemented with 10% FBS and 1% PS was added after 1 hour of incubation. Media was changed to DMEM-Glutamax 1% PS without serum the day after (D0). Cells were fixed and analyzed 5 or 6 days after serum starvation (D5 or D6). Cells expressing the reporter were considered as co-transfected.

### 3D#modelling

Imaris 9.8.0 software was used for 3D modelling.

### Data analysis

All analyses were performed on images of the rostro-dorsal part of the lateral walls of the lateral ventricles to maximize reproducibility.

#### Measurement of phalloidin or YL1/2 fluorescence intensity

Total mean actin (phalloidin fluorescence) or Tyr-tubulin (YL1/2 staining) intensity of fluorescence within cell borders were measured with ImageJ software on the sum projection of the 7 most apical z-stack sections of the cells. Quantifications were normalised with respect to the mean signal intensity in the three closest non-transfected progenitor cells or the mean intensity of progenitor cells of the same image. The results are expressed as the ratio of the values of the cells over the mean values of the progenitor cells.

#### Characterisation of nuclear shape

Nuclei of cells co-transfected with mb-cherry were acquired for their Lamin-B1 staining. Analyses were performed with the Imaris software. For *in vivo* samples, a first loose manual segmentation was performed to exclude nuclei of the untransfected cells. This step was not required for cells *in vitro*. Then an automatic segmentation was performed using Surface and 0.2 grain size resolution. The volume is a quantification of how much space the Surface object occupies (Imaris reference manual). To obtain the pole to pole elongation index, the nucleus is fitted to an ellipse, the dimension of the three principal axes (a, b, c) of the best-fitted ellipsoid into the surface, the moments of inertia along the axes of ellipsoid are computed and optimized to match the moment of inertia of the surface. The ellipsoid has homogeneous mass distribution within the surface object. The pole to pole elongation index is the ellipsoid prolate; define as e_prolate_= (2a^2^/a^2^+b^2^)*(1-((a^2^+b^2^)/2c^2^)) (Imaris reference manual). The Centrin or FOP staining were segmented using dot and 0.25 μm size resolution. The distance of the nucleus to the surface was calculated as the difference in z position of the highest centriole (anchored to the plasma membrane with primary or motile cilia) and the centre of the nucleus. Centre and not the highest part of the nucleus were taken into account, as they are more representative of the nucleus position in a very crowded environment.

#### Characterisation of cell differentiation stages

Only transfected cells with an apical contact at the surface of the ventricle were taken in account. When cells were stained with FOP or Centrin2-GFP, progenitors were considered to have only 2 dots, differentiating cells unresolved or donut like structures, multiciliated cells several resolved dots. Centrin2-GFP accumulate in differentiating cell prior to FOP accumulation and might thus resolved earlier differentiating cells than FOP (not shown).

Sas6 or Deup1 staining, respectively pro-centrioles or deuterosomes, were used to confirm the differentiating cells when possible with the antibodies combinations and fixations.

To increase reproducibility, we counted the stages of the cells surrounding the transfected cells. At P15, images with 60 to 80% (average 65.8%±1.7sem) multiciliated cells were taken in account. At P5, images with 60 to 70% (average 65.6% ±0.4sem) progenitors cells were taken in account.

#### Measurement of H3K27m3, H3K9me2 and phospho-Lamin-A levels *in vivo*

Total mean of H3K27me3 or H3K9me2 or phospho-Lamin-A intensity of fluorescence were measured with ImageJ software on the median plane of the nucleus of cells expressing membrane-Cherry. The measurements were normalized with respect to the mean signal intensity in three closest untransfected progenitor cells.

#### Measurement of NESPRIN2 levels in culture

Total mean of NESPRIN2 intensity of fluorescence were measured with ImageJ software in differentiating ependymal cells *in vitro* on maximum intensity projection of the cell nuclei. Segmentations of the nuclei were performed using automatic thresholding with a triangle filter on Hoechst staining. Background taken in three independent zones in the images were subtracted from the mean value of NESPRIN2 levels for all measurements. The quantification was normalised with respect to the mean signal intensity in the non-transfected progenitor cells or the mean intensity of progenitor cells of the same image. The results are expressed as the ratio of the values of the cells over the mean values of the progenitor cells.

#### Measurement of NESPRIN1 levels in culture

NESPRIN1 is localized both at the centrosome and at the nuclear envelope. At the centrosome, NESPRIN1 levels were quantified with ImageJ software by using a circle of 3.8 μm^2^ centred on the centrosome on maximum intensity projection images. To account for background, the minimum intensity pixel in this area was subtracted from the mean. The levels were normalized with the mean intensity at the centrosome of untransfected progenitor cells in the same image. For nuclear quantification, as centrosome were often located on top of the nucleus, we decided to quantify the nuclear envelope on the nucleus middle plan with ImageJ software by outlining it with a 2 pixel size line. Minimal pixel was subtracted from the mean to take in account the background. Intensity were normalized with respect to the mean signal intensity in the untransfected progenitor cells.

#### Measurement of YAP/TAZ staining

For nuclear quantification, total mean intensity value of the YAP/TAZ staining were measured on the nucleus middle plan with ImageJ software by manual segmentation of the nucleus on the Hoechst channel. The quantification was normalised with respect to the mean signal intensity in the non-transfected progenitor cells or the mean intensity of progenitor cells of the same image. The results are expressed as the ratio of the values of the cells over the mean values of the progenitor cells.

For apical quantification, total mean intensity value of the YAP/TAZ staining were measured on the apical contact plane containing the centrioles with ImageJ software by manual segmentation The quantification was normalised with respect to the mean signal intensity in the non-transfected progenitor cells or the mean intensity of progenitor cells of the same image. The results are expressed as the ratio of the values of the cells over the mean values of the progenitor cells.

#### Measurement of phospho-RB1 staining

Total mean of phospho-RB1 intensity of fluorescence were measured with ImageJ software in differentiating ependymal cells at P5 transfected with MbCherry, on the sum projection of the 5 most central z-stack sections of the nucleus normalised with respect to the mean signal intensity in the three closest untransfected progenitor cells. The results are expressed as the ratio of the values of the transfected cells over the mean values of the three surrounding control cells.

### Statistics

Means of two conditions were compared with the Mann-Whitney test. Means of more than two conditions were analysed by one-way ANOVA followed by Dunn’s multiple comparison tests (GraphPad Prism 6.0). Analyses of cell distribution at different stages were performed by two-way ANOVA with Sidak’s multiple comparisons test. Scatter plots are presented as the mean±sem. ROUT methods were used to remove outliers generated by phalloidin and YAP/TAZ quantification, with a Q value of 1% (GraphPad Prism 6.0). All results were obtained in at least three independent experiments on at least three different animals or cell cultures.

---

**Figure S1:**
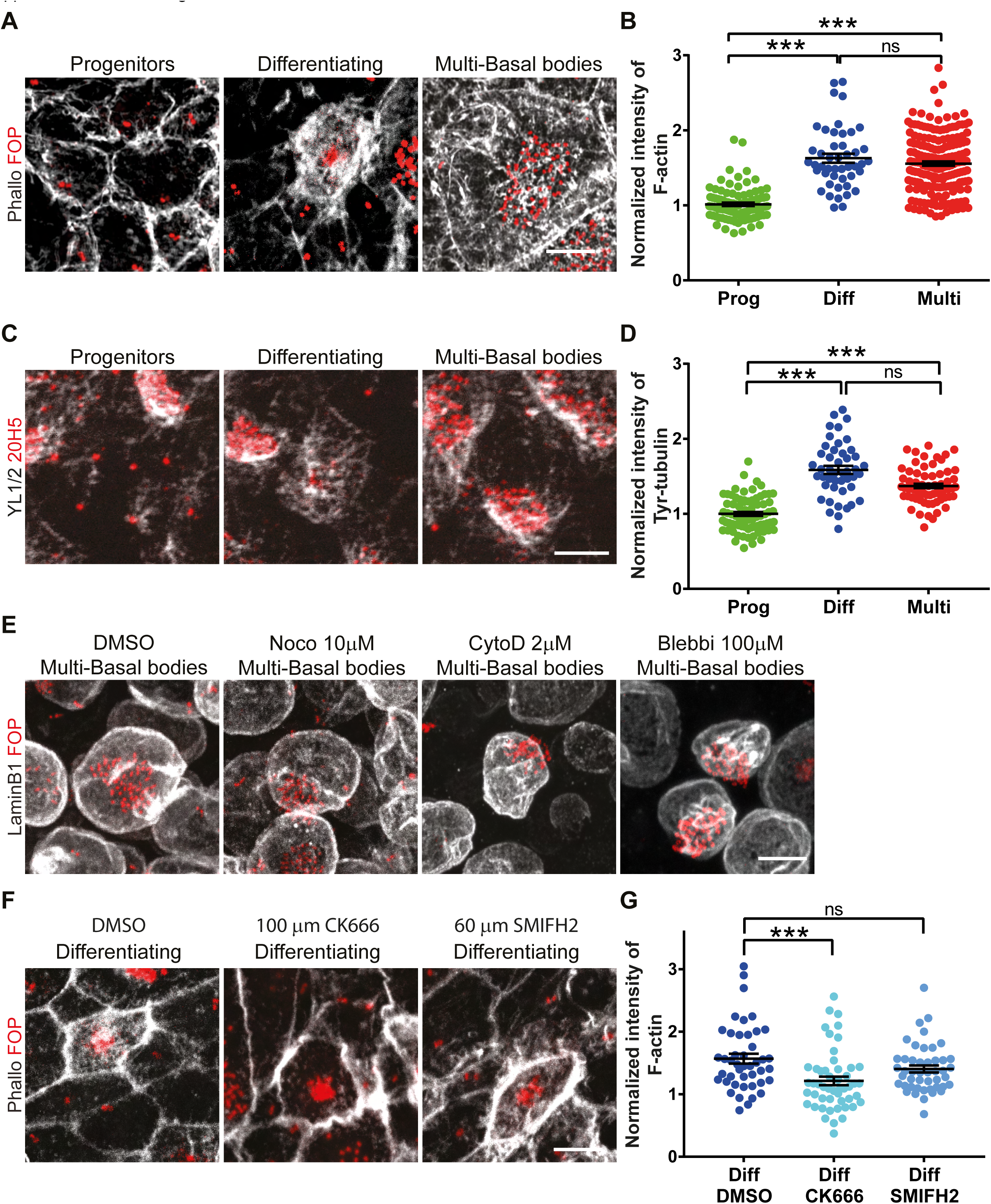
Actin filaments polymerized by the Arp2/3 complex and microtubules accumulate during ependymal differentiation. All staining were performed on whole mount lateral wall during ependymal differentiation (post-natal day 5, P5). The different stages of differentiation are recognized according to centriole staining: 2 dots = Progenitors, 2 dots + clouds or rings = differentiating progenitors, several dots (more than 5) = multi-basal bodies. **A)** Staining of F-actin (Phalloidin, Gray) and centrioles (FOP, Red). **B)** Quantification of the F-actin fluorescence intensity on images as in **A** normalized to the mean fluorescence intensity of F-actin in progenitors cells. Error bars represent the sem of n=96 progenitors, 46 differentiating and 183 multi-basal bodies cells from four mice; p-values from one-way ANOVA followed by Dunn’s multiple comparison test; *** P≤0.001; ns P>0.05. **C)** Staining of Tyrosinated-Tubulin (YL1/2= polymerizing tubulin, Gray) and Centrin (20H5= centrioles, Red). **D)** Quantification of the Tyr-tubulin fluorescence intensity on images as in **C** normalized to the mean fluorescence intensity of Tyr-tubulin in progenitors. Error bars represent the sem of n=75 progenitors, 46 differentiating and 66 multi-basal body cells from three mice; p-values from one-way ANOVA followed by Dunn’s multiple comparison test; *** P≤0.001; ns P>0.05. **E)** Staining of lamin (Lamin-B1, Gray) and centrioles (FOP, Red) on explant treated with DMSO or nocodazole (microtubule inhibitor 10 μM) or cytochalasin-D (F-actin inhibitor, 2 μM) or Blebbistatin (100 μM, Myosin II inhibitor).for 20 hours. **F)** Staining of F-actin (Phalloidin, Gray) and centrioles (FOP, Red) on explant at P4 or P5 treated for 20 hours with DMSO or CK666 (100 μM, Arp2/3 complex inhibitor) or SMIFH2 (60 μM, formin inhibitor). **G)** Quantification of the F-actin fluorescence intensity on differentiating ependymal cells on images as in **F** normalized to the mean fluorescence intensity of F-actin in progenitors of relative images. Error bars represent the sem of n=44, 46 and 50 differentiating cells treated with DMSO, CK666 or SMIFH2 respectively from four independent mice; p-values from one-way ANOVA followed by Dunn’s multiple comparison test; *** P≤0.001; ns P>0.05. Scale bar = 5 μm.

**Figure S2:**
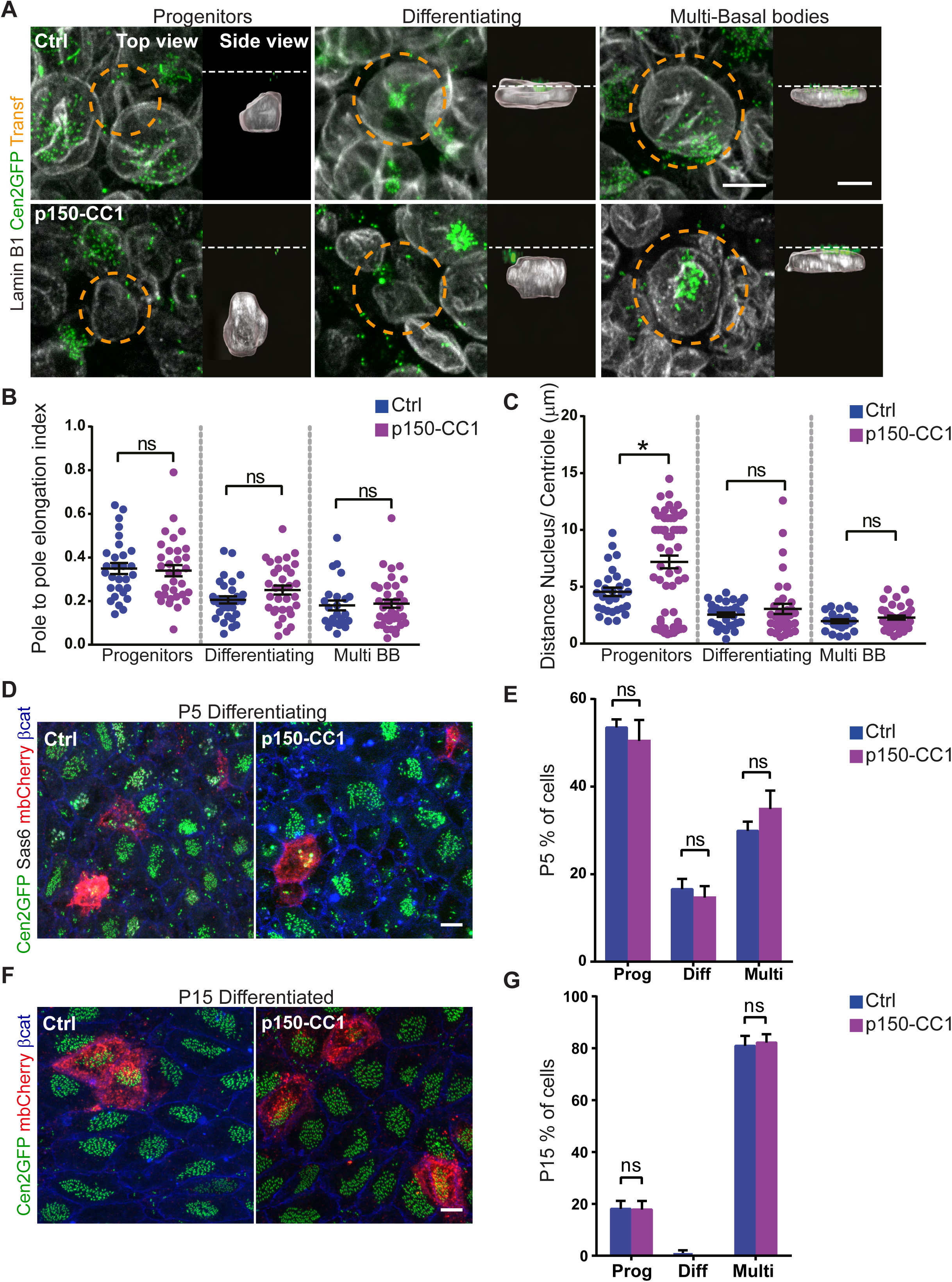
Disturbing Dynein does not affect ependymal differentiation. Wild type or Centrin2-GFP mice transfected at P1 with mb-cherry as a reporter and either a control plasmid (Ctrl) or the p150-CC1 domain (dominant negative of dynein, p150-CC1). **A)** Staining of Lamin-B1 (nuclear lamina, Gray) and Centrin 2-GFP (Cen2GFP, centrioles, Green) during ependymal cell differentiation (P5) on whole mount lateral wall. Transfected cells, either Ctrl (top row) or p150-CC1 (bottom row), are circled with orange dashed lines. Top view is a maximum projection of z-stacks. Side view is an Imaris 3D projection with segmented nuclei. The white dashed lines represent the plasma membrane. **(B, C)** Quantification of the nuclear shape during the different steps of ependymal differentiation on Imaris segmented nuclei as in **A**. **B)** Pole to pole elongation index. **C)** Distance between the center of the nucleus and the highest centriole. Error bars represent the sem of n=29/ 59 progenitors, 30/ 35 differentiating and 23/ 36 multi-basal bodies (Multi BB) transfected cells (Ctrl/ p150-CC1) from three mice; p-values from Mann-Whitney test; ns, P>0.05. * p=0.0132. Control values are the same than on Figure 4. **D)** Staining of Centrin2-GFP (Cen2GFP, centrioles, Green), Sas6 (procentrioles, Gray), mb-cherry (reporter, Red) and beta-catenin (junction, Blue) during ependymal cell differentiation (P5) on whole mount lateral wall. **E)** Quantification of the percentage of transfected cells as in **D** at each stages of differentiation at P5. (Prog=progenitors, Diff=Differentiating, Multi=multi-basal bodies) Error bars represent the sem of 5 experiments n=174 (Ctrl) and 175 (p150-CC1) transfected cells; p-values from two-way ANOVA followed by Sidak’s multiple comparisons test; ns, P>0.05. Control values are the same than on Figure 2 as the experiments were performed at the same time. **F)** Staining of Centrin2-GFP (Cen2GFP, centrioles, Green), mb-cherry (reporter, Red) and beta-catenin (junction, Blue) in differentiated cells (P15) on whole mount lateral wall. **G)** Quantification of the percentage of transfected cells as in **F** remaining as progenitors (Prog), still differentiating (Diff) or differentiated in ependymal cells (Multi) at P15. Error bars represent the sem of 4 (Ctrl) or 4 (p150-CC1) experiments n=83 (Ctrl) and 65 (p150-CC1) transfected cells; p-values from two-way ANOVA followed by Sidak’s multiple comparisons test; *** P≤0.0001; ns, P>0.05. Control values are the same than on Figure 2. Scale bar = 5 μm.

**Figure S3:**
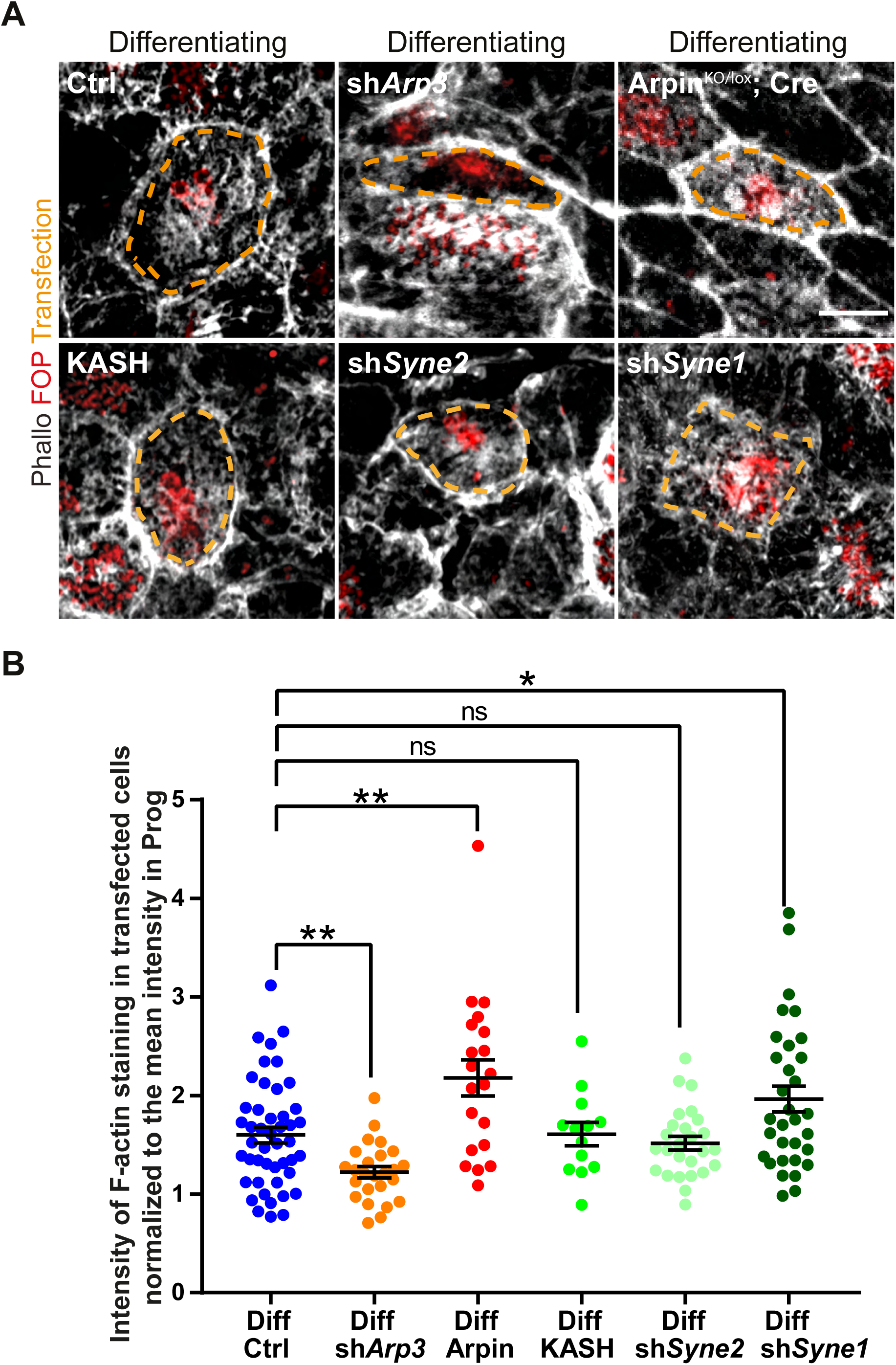
Effects of tampering with actin or the LINC complex on actin organization in differentiating cells. **A**) Staining of F-actin (Phalloidin, Gray) and centrioles (FOP, Red) on differentiating cells transfected with mb-cherry (reporter, orange dashed line) and either a control plasmid (Ctrl) or an shRNA against *Arp3* (sh*Arp3*) or the KASH domain or an shRNA against *Syne2* or against *Syne1* or Arpin^KO/lox^ mice electroporated with a plasmid encoding for the Cre recombinase. **B)** Quantification of the F-actin fluorescence intensity on images as in **A** normalized to the mean fluorescence intensity of F-actin in un-transfected progenitors of relative images. Error bars represent the sem of n=46, 26, 20, 13, 26, 32 transfected differentiating cells (Ctrl (wild type or Arpin^lox/lox^ mice) or sh*Arp3* or Cre (Arpin^KO/lox^ mice expressing the Cre recombinase) or OE KASH or sh*Syne2* or shSyne1 respectively) from four mice; p-values from one-way ANOVA followed by Dunn’s multiple comparison test; Ctrl vs sh*Arp3* **p=0.0076; Ctrl vs Arpin^KO/lox^; Cre **p=0.0053; Ctrl vs shSyne1 * p=0.04. Scale bar = 5 μm.

**Figure S4:**
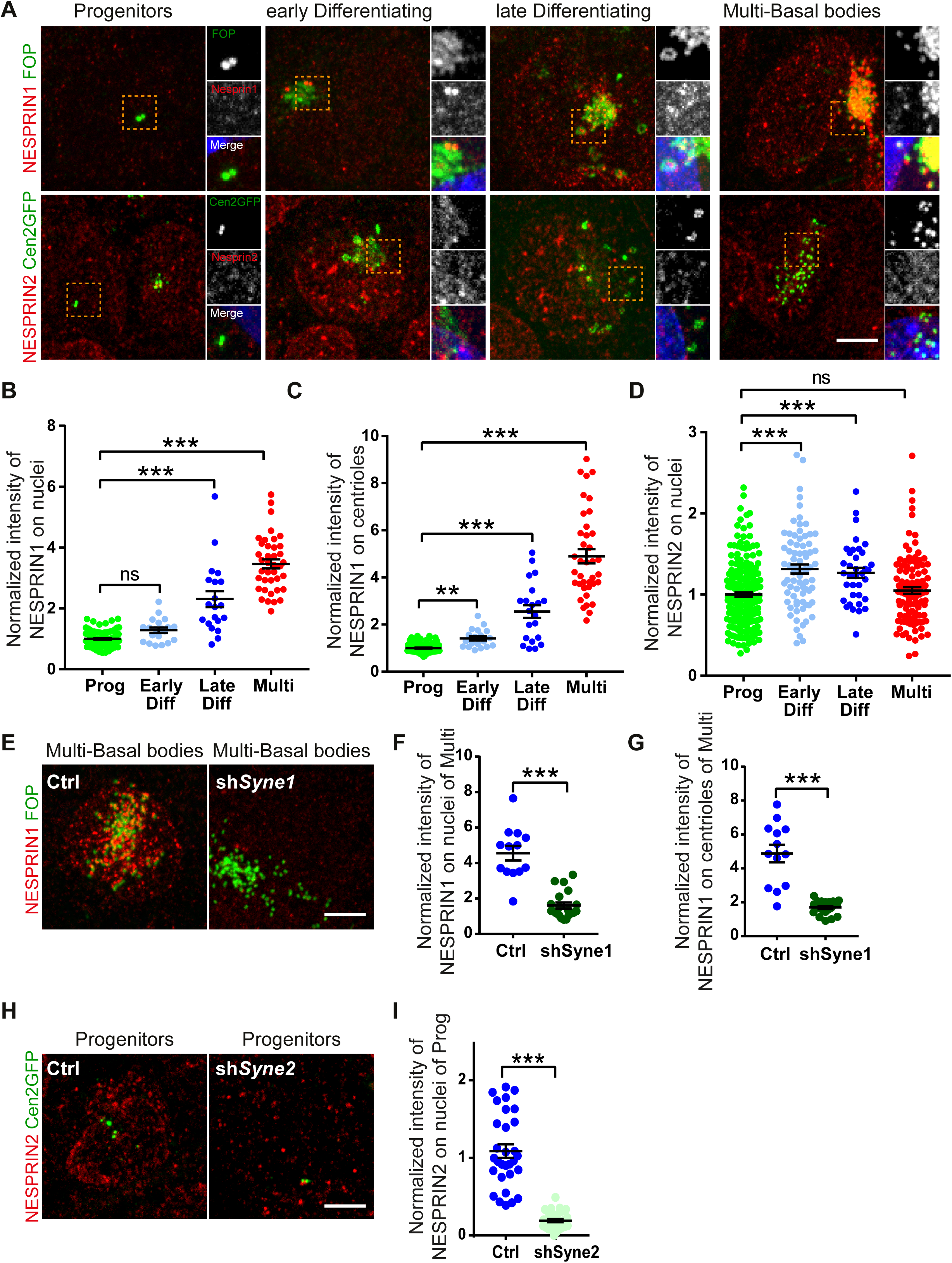
NESPRIN1 and 2 localization and shRNA validation. Related to Figure 5. **A)** Staining of NESPRIN1 (Top, Red) or NESPRIN2 (Bottom, Red) and a centriolar marker either FOP (Top, Green) or Centrin 2-GFP (Bottom, Cen2GFP, Green) during ependymal cell differentiation *in vitro* (6 days after serum withdrawal). Small insets are 2x single channel zoom images of the orange dashed squares. Blue is Hoechst staining. **(B-D)** Quantification of the intensity of fluorescence of either NESPRIN1 staining at the centrosome (**B**) or at the nuclear envelope (**C**) or of NESPRIN2 staining at the nuclear envelope (**D**) at each stages of differentiation: progenitor (Prog), differentiating (Diff either early (centriolar marker as a cloud) or late (centriolar marker as ring structures)) or already differentiated (Multi) on *in vitro* culture as in **A**, normalized to the mean intensity of progenitor cells for each staining on each images. Error bars represent the sem of n=102 prog, 20 early diff, 20 late diff and 39 multi cells for NESPRIN1 quantification; n=217 prog, 75 early diff, 36 late diff and 107 multi cells for NESPRIN2 quantification; from three independent cultures; p-values from Mann-Whitney test *** P≤0.0001; ns, P>0.05 **(E-I)** Cells were transfected *in vitro* at D0 with shRNA (either Ctrl or against *Syne1* or *Syne2*) and tomato (as a reporter) and fixed 6 days after differentiation onset (=serum withdrawal). **E)** Staining of NESPRIN1 (LINC complex, Red) in transfected multi basal body cells either Ctrl or depleted for NESPRIN1 (sh*Syne1*). **(F-G)** Quantification of the intensity of fluorescence of NESPRIN1 staining *in vitro* at the centrosome **(F)** or at the nuclear envelope **(G)** in transfected multi basal body cells at D6 either control (Ctrl) or depleted for NESPRIN1 (sh*Syne1*) as in **E,** normalized to the mean of intensity of untransfected progenitor cells on each images. Error bars represent the sem of n=13 and 20 cells for Ctrl or sh*Syne1* respectively; from three independent cultures; p-values from Mann-Whitney test *** P≤0.0001; **H)** Staining of NESPRIN2 (LINC complex, Red) in transfected progenitor cells either Ctrl or depleted for NESPRIN2 (sh*Syne2*). **I)** Quantification of the intensity of fluorescence of NESPRIN2 staining *in vitro* in progenitor cells at D6 either control (Ctrl) or depleted for NESPRIN2 (sh*Syne2*) as in **H,** normalized to the mean of intensity of untransfected progenitor cells on each images. Error bars represent the sem of n=30 cells for Ctrl or sh*Syne2*; from three independent cultures; p-values from Mann-Whitney test *** P≤0.0001; Scale bar = 5 μm.

**Figure S5:**
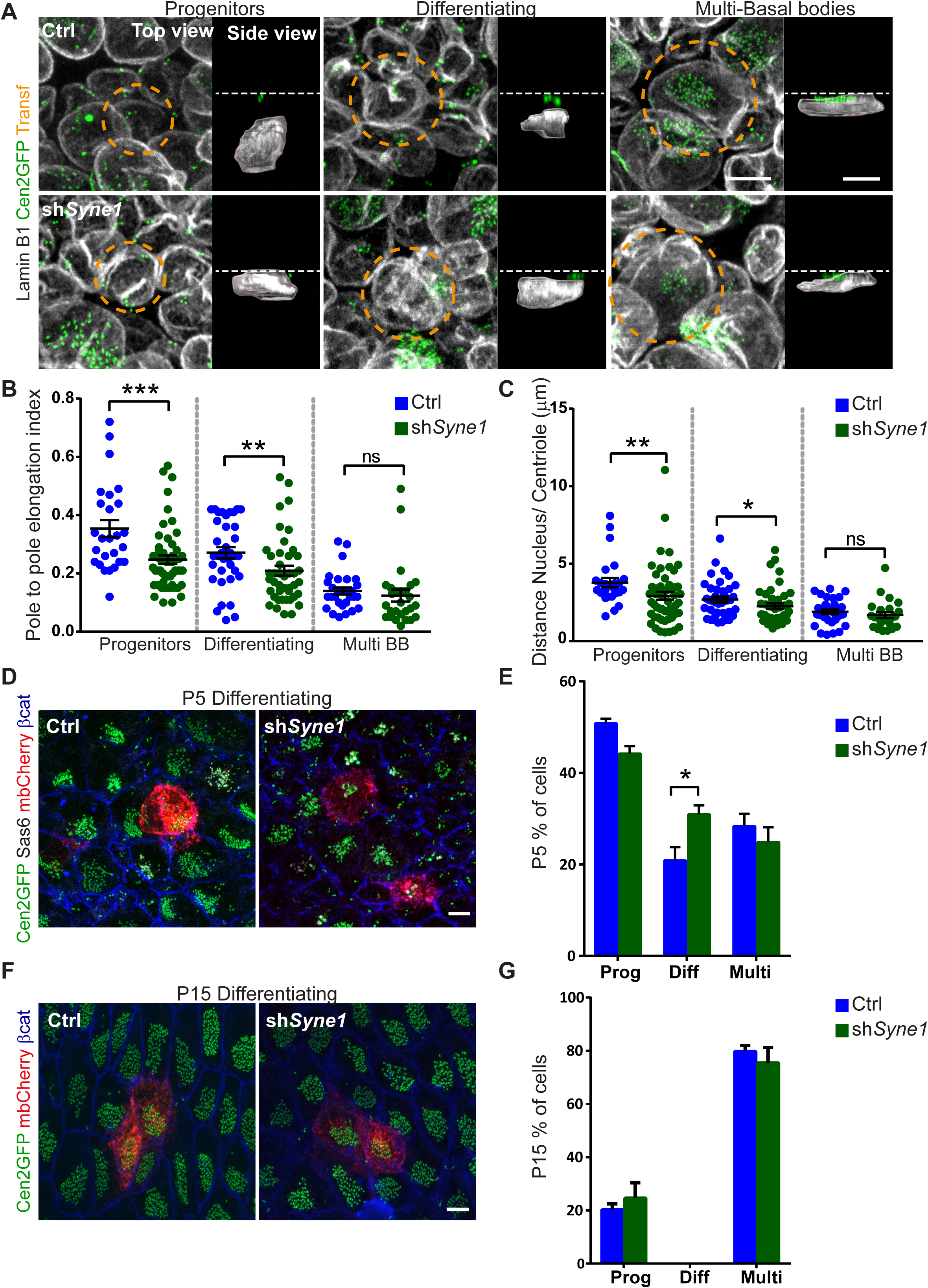
Decreasing NESPRIN1 advances nuclear re-shaping and ependymal differentiation. Wild type or Centrin2-GFP mice transfected at P1 with mb-cherry as a reporter and either a control plasmid (Ctrl) or an shRNA against *Syne1* (sh*Syne1* to deplete NESPRIN1). **A)** Staining of Lamin-B1 (nuclear lamina, Gray) and Centrin 2-GFP (Cen2GFP, centrioles, Green) during ependymal cell differentiation (P5) on whole mount lateral wall. Transfected cells, either Ctrl (top row) or sh*Syne1* (bottom row), are circled with orange dashed lines. Top view is a maximum projection of z-stacks. Side view is an Imaris 3D projection with segmented nuclei. The white dashed lines represent the plasma membrane. **(B, C)** Quantification of the nuclear shape during the different steps of ependymal differentiation on Imaris segmented nuclei as in **A**. **B)** Pole to pole elongation index. **C)** Distance between the center of the nucleus and the highest centriole. Error bars represent the sem of n=26/ 54 progenitors, 40/ 42 differentiating and 24/ 30 multi-basal bodies (Multi BB) transfected cells for Ctrl or sh*Syne1* respectively from four mice; p-values from Mann-Whitney test; ns, P>0.05;* p=0.043; *** P≤0.001. Control values are the same than on Figure 2. **D)** Staining of Centrin2-GFP (Cen2GFP, centrioles, Green), mb-cherry (reporter, Red) and beta-catenin (junction, Blue) in differentiated cells (P5) on whole mount lateral wall. **E)** Quantification of the percentage of transfected cells as in **D** differentiated in ependymal cells (Multi), still differentiating (Diff) or remaining as progenitors (Prog) at P5. Error bars represent the sem of 6 (Ctrl) or 5 (sh*Syne1*) experiments n=148 (Ctrl) and 168 (sh*Syne1*) transfected cells; p-values from two-way ANOVA followed by Sidak’s multiple comparisons test; * P=0.031; ns. Control values are the same than on Figure 4. **F)** Staining of Centrin2-GFP (Cen2GFP, centrioles, Green), mb-cherry (reporter, Red) and beta-catenin (junction, Blue) in differentiated cells (P15) on whole mount lateral wall. **G)** Quantification of the percentage of transfected cells as in **F** remaining as progenitors (Prog), still differentiating (Diff) or differentiated in ependymal cells (Multi) at P15. Error bars represent the sem of 6 (Ctrl) or 2 (sh*Syne1*) experiments n=148 (Ctrl) and 39 (sh*Syne1*) transfected cells; p-values from two-way ANOVA followed by Sidak’s multiple comparisons test; ns, P>0.05. Control values are the same than on Figure 4. Scale bar = 5 μm.

**Figure S6:**
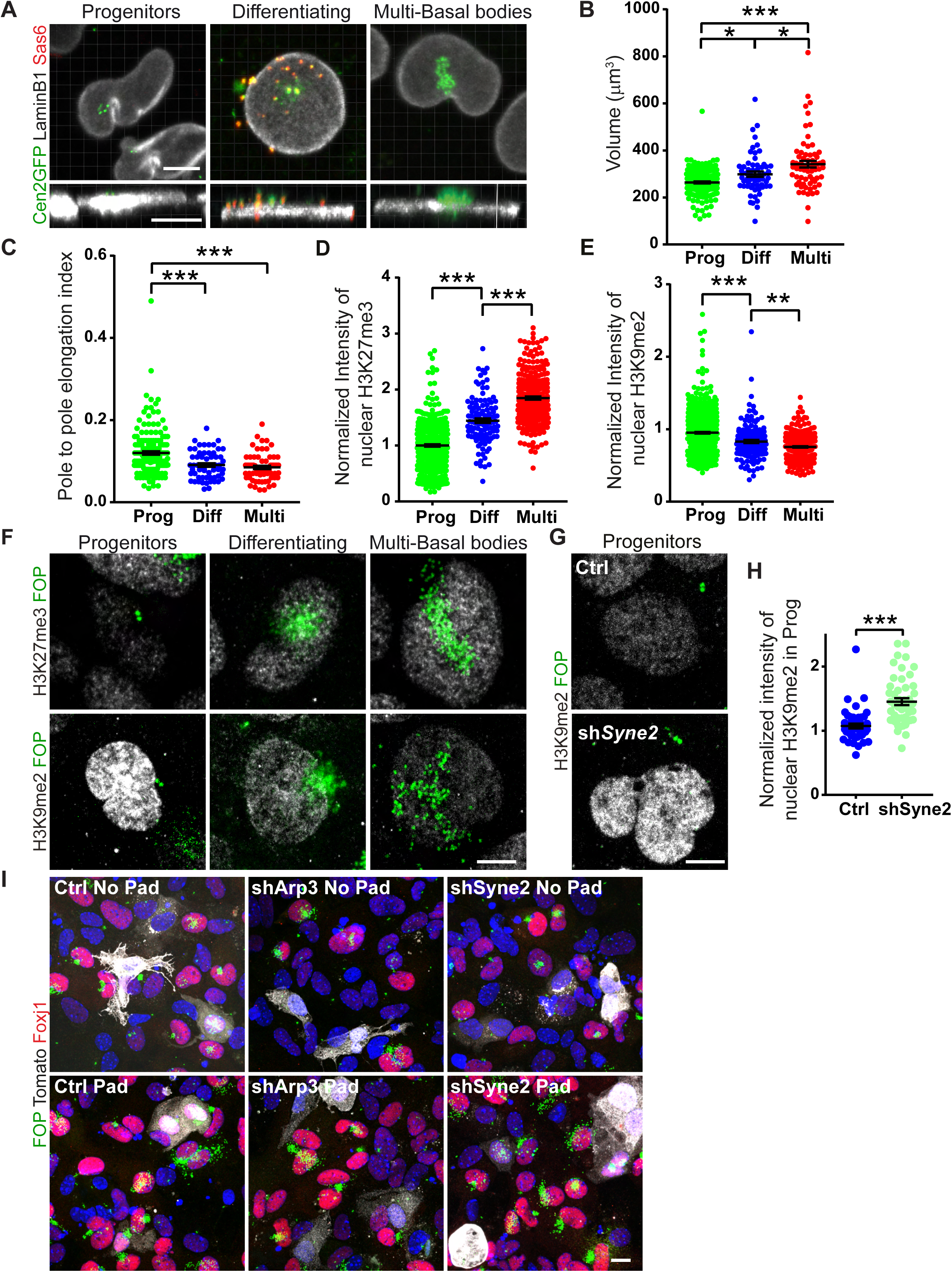
Validation of the role of nuclear deformation in *in vitro* culture. Related to Figure 6. **A)** Staining of Lamina (Lamin-B1, Gray), procentrioles (Sas6, Red) and centrioles (Cen2GFP, Green) during ependymal cell differentiation *in vitro* (5 days after serum removal, D5). The different stages of differentiation are recognized according to the Centrin2-GFP and Sas6 pattern: 2 dots of Centrin + no Sas6 = Progenitors, 2 dots + clouds or rings of Centrin + dots of Sas6 = differentiating progenitors, several dots of Centrin (more than 5) and no Sas6 = multi-basal bodies. Top and side views are Imaris 3D projection of a z-stack. **(B-C)** Quantification of nuclear shapes during the different steps of ependymal differentiation *in vitro* on Imaris segmented nuclei. **B)** Volume. **C)** Pole to pole elongation index. Error bars represent the sem of n=174 progenitors (Prog), 60 differentiating (Diff) and 67 multi-basal body (Multi) cells from 5 different cultures. P-values from one-way ANOVA followed by Dunn’s multiple comparison test; for the volume; * p=0.0129 (Prog vs Diff); * p=0.0396 (Diff vs Multi); *** P≤0.001. **(D, E)** Quantification of the fluorescence intensity of H3K27me3 (**D**) or H3K9me2 (**E**) staining in cells 6 days after differentiation onset at different stages of differentiation (Prog= progenitors, Diff= differentiating, Multi= multiciliated differentiated cells) on images as in **F** normalized to the mean of intensity in progenitors of the same image. Error bar represent the sem of n=407, 111 and 275 cells for Prog, Diff, Multi cells respectively in H3K27me3 staining and n=681, 143 and 338 cells for Prog, Diff, Multi respectively in H3K9me2 staining in three independent experiments each. p-values were determined by one-way ANOVA followed by Dunn’s multiple comparison test; *** P≤0.001; ** P<0.01. **F)** Staining of H3K27me3 (Gray, top panel) or H3K9me2 (Gray, bottom panel) and FOP (centrioles, Green) during ependymal differentiation (P5). **(G, H)** Cells were transfected at differentiation onset (D0) with Tomato as a reporter and either a control plasmid (Ctrl) or an shRNA against *Syne2* (sh*Syne2*) and fixed 6 days after serum withdrawal. **G)** Staining of centrioles (FOP, Green) and H3K9me2 (Gray) in cells either control (Ctrl) or depleted for NESPRIN2 (sh*Syne2*). **H)** Quantification of the levels of H3K9me2 fluorescence intensity in Ctrl or sh*Syne2* cells. Error bar represent the sem of n=47 or 49 cells for Ctrl and sh*Syne2* respectively in two independent experiments. p-values from Mann-Whitney test; *** P≤0.0001. **I)** Progenitor cells *in vitro* were transfected with shRNA either control (left panel) or against *Arp3* (middle panel) or against *Syne2* (right panel) at differentiation onset (D0= serum removal). They were then confined (Pad) or not (No Pad) under an agarose pad for 6 days, fixed and stained with FOP (centriole, Green), Foxj1 (transcription factor specific of multiciliated differentiating and differentiated cells, Red), Tomato (reporter, Gray) and Hoechst (DNA, Blue). Scale bar = 5 μm.

**Figure S7:**
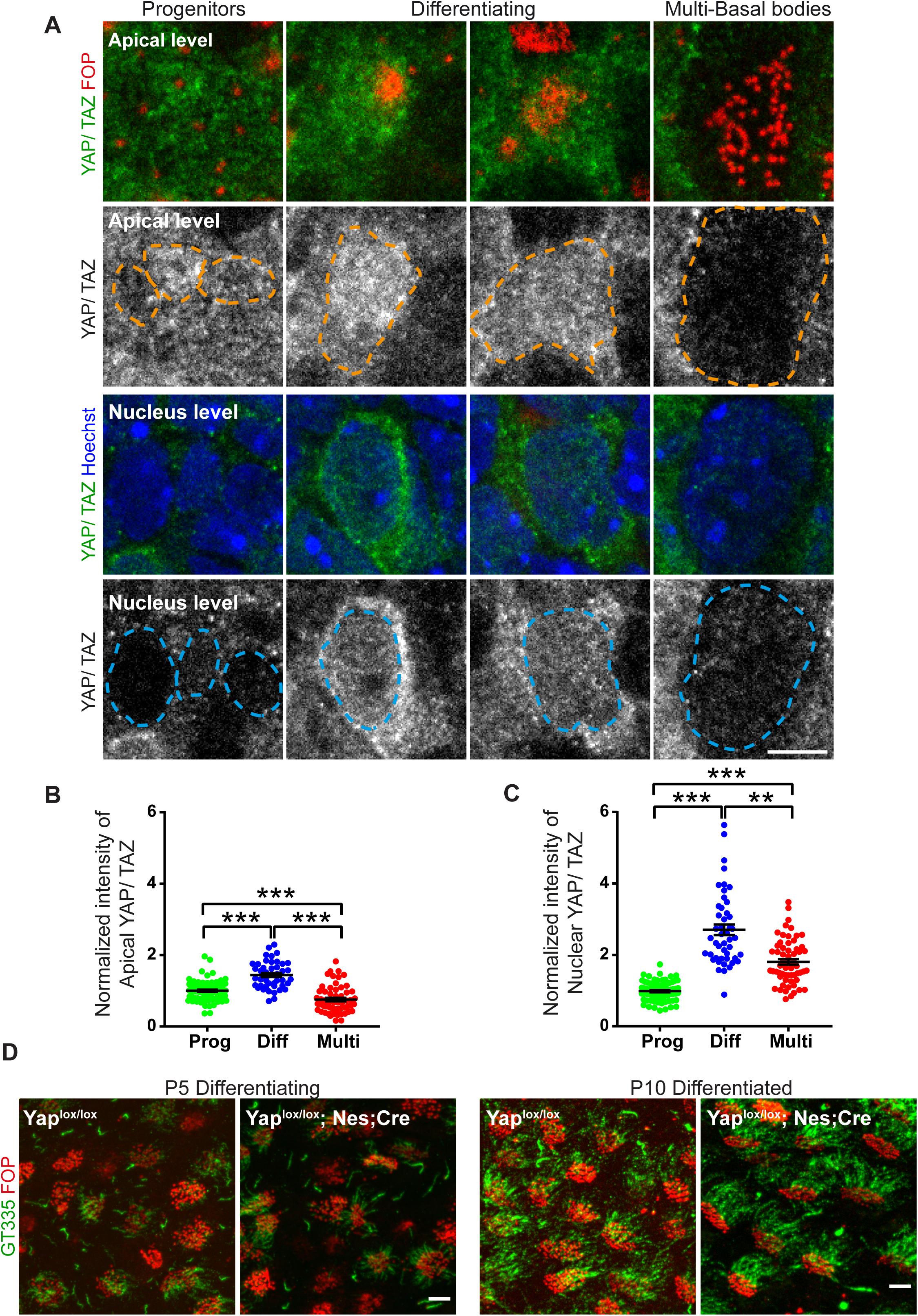
Role of YAP in ependymal differentiation. **A**) Staining of YAP/ TAZ (Green on the merge images or Gray on the single channel images), centrioles (FOP, Red) or nucleus (Hoechst, Blue) during ependymal cell differentiation on whole mount lateral wall at P5. Top row is the z-stack of the most apical plans (excluding the nucleus, Apical level). Bottom row is the z-stack of the plans containing the nucleus (Nucleus level).The different stages of differentiation are recognized according to the FOP pattern: 2 dots = Progenitors, 2 dots + clouds or rings = differentiating progenitors, several dots (more than 5) = multi-basal bodies. **B)** Quantification of the fluorescence intensity of YAP/TAZ staining at the apical level in cells at the different steps of ependymal differentiation at P5, normalized to the mean intensity of progenitor cells. Error bars represent the sem of n=40 progenitors, 37 differentiating and 29 multi-basal bodies cells from 2 mices p-values were determined by one-way ANOVA followed by Dunn’s multiple comparison test; *** P≤0.001. **C)** Quantification of the fluorescence intensity of YAP/TAZ staining at the nucleus level in cells at the different steps of ependymal differentiation at P5, normalized to the mean intensity of progenitor cells. Error bars represent the sem of n=40 progenitors, 37 differentiating and 29 multi-basal bodies cells from 2 mices p-values were determined by one-way ANOVA followed by Dunn’s multiple comparison test; *** P≤0.001. **D)** Staining of whole mount lateral walls of P5 or P10 animals either Control (YAP^lox/lox^) or mutant for YAP (YAP^lox/lox^; Nes:Cre) stained for centrioles (FOP, Red) and cilia (GT335 polyglutamylated tubulin, Green) Scale bar = 5 μm.

**Figure S8:**
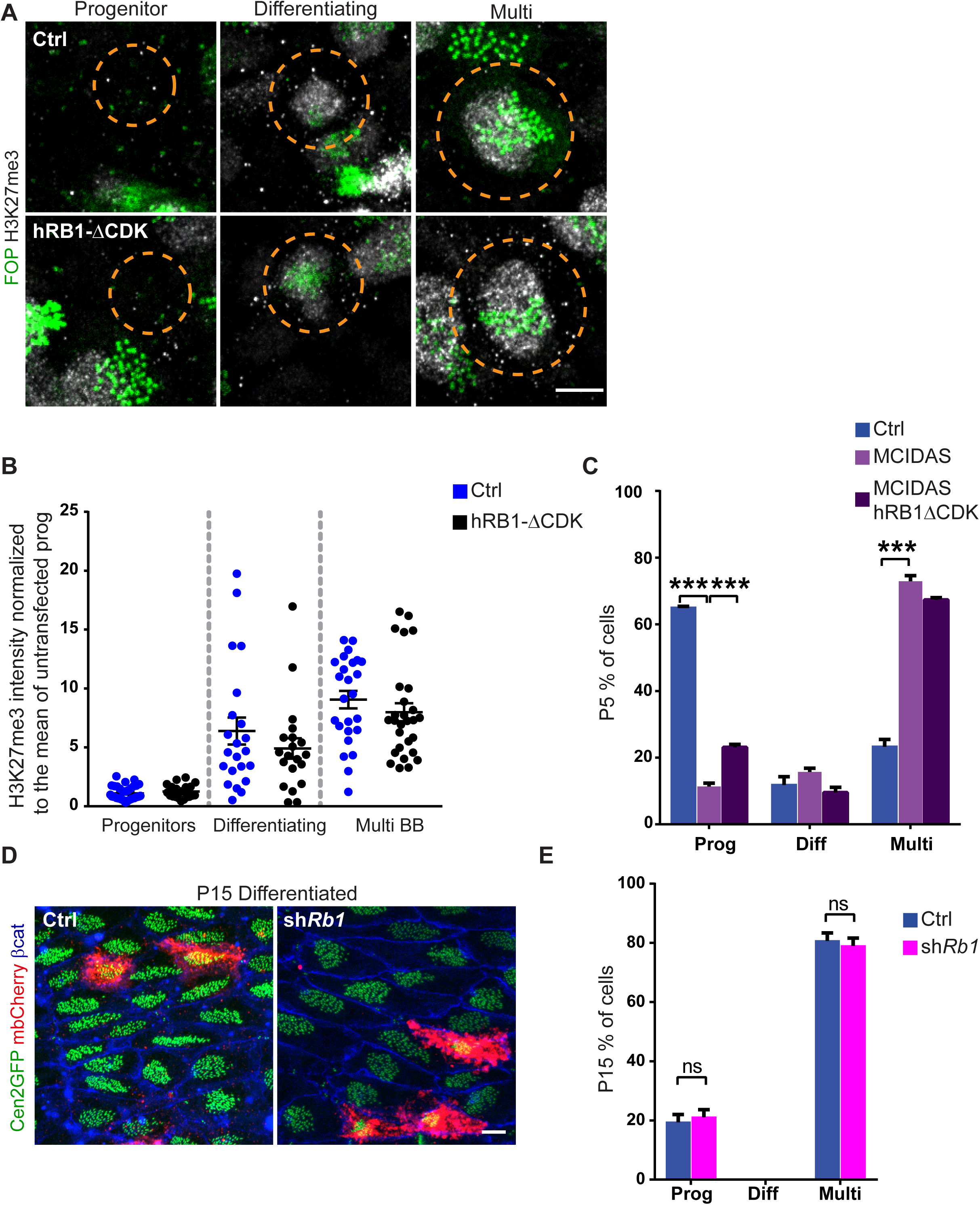
H3K27 is methylated in presence of hRB1-ΔCDK, while depleting RB1 does not affect ependymal differentiation. Related to Figure 7. **A)** Staining of centriole (FOP, Green) and H3K27me3 (Gray) differentiating ependymal cells (P5) on whole mount lateral wall of wild-type animals ransfected with mb-cherry (Orange dashed line), as a reporter and either a control plasmid or hRB1-ΔCDK. **B)** Quantification of the fluorescence intensity of H3K27me3 staining in transfected cells as in **A** normalized to the mean intensity of three surrounding untransfected progenitor cells. Error bar represent the sem of n=35/ 20Prog, 22/ 20 Diff, and 25/ 28 Multi cells for Ctrl/ OE hRB1-ΔCDK respectively in two animals. p-values from Mann-Whitney test none significant (P>0.05) comparing Ctrl and hRB1-ΔCDK for each stages. **C)** Quantification of the percentage of transfected cells at P5 on whole mount lateral wall of wild-type mice transfected at P1 with control plasmids (Ctrl= shNeg-IRES-GFP+ shNeg) or MCIDAS (shNeg-IRES-GFP+ MCIDAS) or both MCIDAS and unphosphorylatable RB1 (hRB1-ΔCDK –IRES-GFP+ MCIDAS) in the different stages of differentiation: Progenitors (Prog), differentiating (Diff) or differentiated (Multi). Error bars represent the sem of 3 (Ctrl) or 3 (MCIDAS) or 2 (MCIDAS + hRB1-ΔCDK) experiments, n=91 (Ctrl), 98 (MCIDAS) and 43 (MCIDAS + hRB1-ΔCDK) transfected cells; p-values from two-way ANOVA followed by Sidak’s multiple comparisons test; *** P≤0.0001. **D)** Staining of Centrin2-GFP (Cen2GFP, centrioles, Green), mb-cherry (reporter, Red) and beta-catenin (junctions, Blue) in differentiated cells (P15) on whole mount lateral wall of Centrin2-GFP mice transfected at P1 with mb-cherry and either a control plasmid (Ctrl) or sh*Rb1*. **E)** Quantification of the percentage of transfected cells as in **D**, either Control (ctrl) or sh*Rb1* remaining as progenitors (Prog), still differentiating (Diff) or differentiated in ependymal cells (Multi) at P15. Error bars represent the sem of 4 experiments n=146 (Ctrl same as in Figure 6D), 81 (shRb1) transfected cells; p-values from two-way ANOVA followed by Sidak’s multiple comparisons test; ns, P>0.05. Scale bar = 5 μm.

